# Birth of an organelle: molecular mechanism of lipid droplet biogenesis

**DOI:** 10.1101/2023.07.28.550987

**Authors:** Vincent Nieto, Jackson Crowley, Denys Santos, Luca Monticelli

## Abstract

Lipid droplets (LDs) are cellular organelles regulating energy and lipid metabolism. Generated in the endoplasmic reticulum (ER) by phase separation of neutral lipids (e.g., triglycerides), nascent LDs resemble lens-shaped blisters, grow into spherical droplets, and eventually emerge from the ER membrane, generally towards the cytosol – a process known as budding. Images of both nascent and mature LDs are available, but the mechanism of biogenesis has never been observed experimentally.

Here we identify the mechanism of the initial steps in LD biogenesis using computer simulations at the molecular level, emulating LD growth in ER-mimicking membranes. We find that LDs bud towards the cytosol only when sufficient asymmetry is generated between the two membrane leaflets: the budding transition is independent of membrane morphology, lipid composition, and LD size. Seipin, a protein essential for correct LD biogenesis, is *per se* not sufficient to promote budding, but it stabilizes LD-ER contact sites. Localization of triglyceride synthesis in the proximity of seipin is necessary to avoid nucleation of multiple LDs – a possible cause of aberrant phenotypes. In contrast, localization of phospholipid synthesis has no effect on the mechanism of budding. Our new methodology paves the way to simulations of organelle and cell biogenesis.

---

Adenosine triphosphate (ATP) provides the energy to drive most chemical reactions in living organisms^1^, but cannot cannot easily be stored^2^. Lipids, instead, are a convenient way to store energy. Cells package them in lipid droplets (LDs) – crucial hubs of lipid metabolism and central in membrane biogenesis and several non-metabolic processes (e.g., protein quality control and viral infections^3^). Deficiencies in LD formation or functioning result in impaired cell metabolism, cell stresses, and numerous diseases, e.g., lipodystrophies, liver steatosis, type II diabetes, and neurodegeneration^4-6^. LDs are generated mostly in the endoplasmic reticulum (ER), the prime site for lipid synthesis. LD biogenesis starts with the synthesis of neutral lipids (such as triglycerides and sterol esters)^6^. When the concentration of neutral lipids in the ER membrane reaches a certain threshold, they phase-separate from the phospholipids constituting the ER bilayer^7^. Phase separation starts with the nucleation of a lens-shaped nascent LD, sandwiched between the two leaflets of the ER bilayer; the nascent LD grows as more neutral lipids are synthesized, and eventually buds out of the ER, generally towards the cytosol, coated by the cytosolic leaflet of the ER bilayer, i.e., a mono-molecular layer of phospholipids and proteins. Mature LDs generally remain attached to the ER via LD-ER contact sites marked by seipin, an oligomeric protein with an important role in preventing pathological phenotypes^8,9^. However, the mechanism of LD biogenesis has never been observed, due to the fluid nature of LDs and the insufficient spatial and time resolution of current microscopy techniques: observable LDs have a size of hundreds of nanometers or more, but nascent LDs are much smaller, and isolating the different steps of the process is proving a formidable challenge for structural biology^5,6^. In the absence of a high-resolution view of the mechanism of biogenesis, several questions remain open on the driving forces and the molecular factors determining LD budding. For instance, continuum theory predicts that nascent LDs should spontaneously bud off from a symmetric bilayer when their size reaches a few tens of nm^10^, provided a sufficiently low surface tension in the bilayer, but validation of the theory is problematic as it would require fine control over the synthesis of both neutral and polar lipids. Contrasting results have been reported on the role and the nature of ER membrane asymmetry: Choudhary *et al*. proposed that directional budding is determined by intrinsic curvature of ER phospholipids^11^, while Chorley *et al*. proposed that it depends on the asymmetry in surface tension (i.e., surface density) between ER leaflets^12^. The function of seipin is also not completely clear: simulations and experiments suggested that it may trap triglycerides^13-15^, therefore affecting LD nucleation and growth by ripening, but its localization at the LD-ER contact site raises questions on a possible role also in the budding process.

Here we address the questions on the driving forces, the threshold size, and the mechanism of LD budding, as well as on the role of seipin, by simulating the initial steps of LD biogenesis at the molecular level. To this end, we develop a novel protocol that allows molecular simulations with increasing number of particles, and we use it to emulate the synthesis of different lipids in membranes mimicking ER tubules. The outcome is an unprecedented view of the birth of a cellular organelle, with nanosecond time resolution and sub-nanometer spatial resolution. We find that triglyceride synthesis results in budding of stable and defect-free LDs only under specific conditions: low surface tension, active generation of asymmetry between the ER leaflets, presence of seipin, a specific composition of the ER membrane, and a precise regulation of both triglyceride and phospholipid synthesis. Our results provide a consistent interpretation for published experimental data and specific predictions amenable to experimental validation.

## Synthesis of triglycerides *per se* does not induce lipid droplet budding

In cells, LDs generally bud off from tubular regions of the ER^16^, and ER curvature has been proposed to catalyze LD assembly^17^. To mimic biogenesis in the ER, we carried out molecular dynamics (MD) simulations of bilayer membranes with a tubular shape, with size comparable to ER tubules^18,19^ (see Methods) and negligible surface tension (SI.4), effectively mimicking biological systems^20^. We then devised an algorithm, coined POP-MD, to emulate lipid synthesis and simulate LD growth, by having triglyceride (TG) molecules pop into the system at user-defined locations and rates (Table 1). As TG concentration reached a critical threshold, initially well-dispersed oil molecules rapidly formed one or more lens-shaped nascent LDs; nucleation occurred on time scales of hundreds of ns. TG synthesis in random locations within the membrane induced nucleation of multiple LDs, which did not coalesce into a single LD within the simulation time (10 μs), due to the long time scales required for LD diffusion (Fig. 1a, Supplementary Movie 1). During LD growth, about 2% of TG remained free in the bilayer, in agreement with previous data^21-23^ and with our own equilibrium simulations, suggesting that LD growth by POP-MD is not far from equilibrium.

**Table 1.**
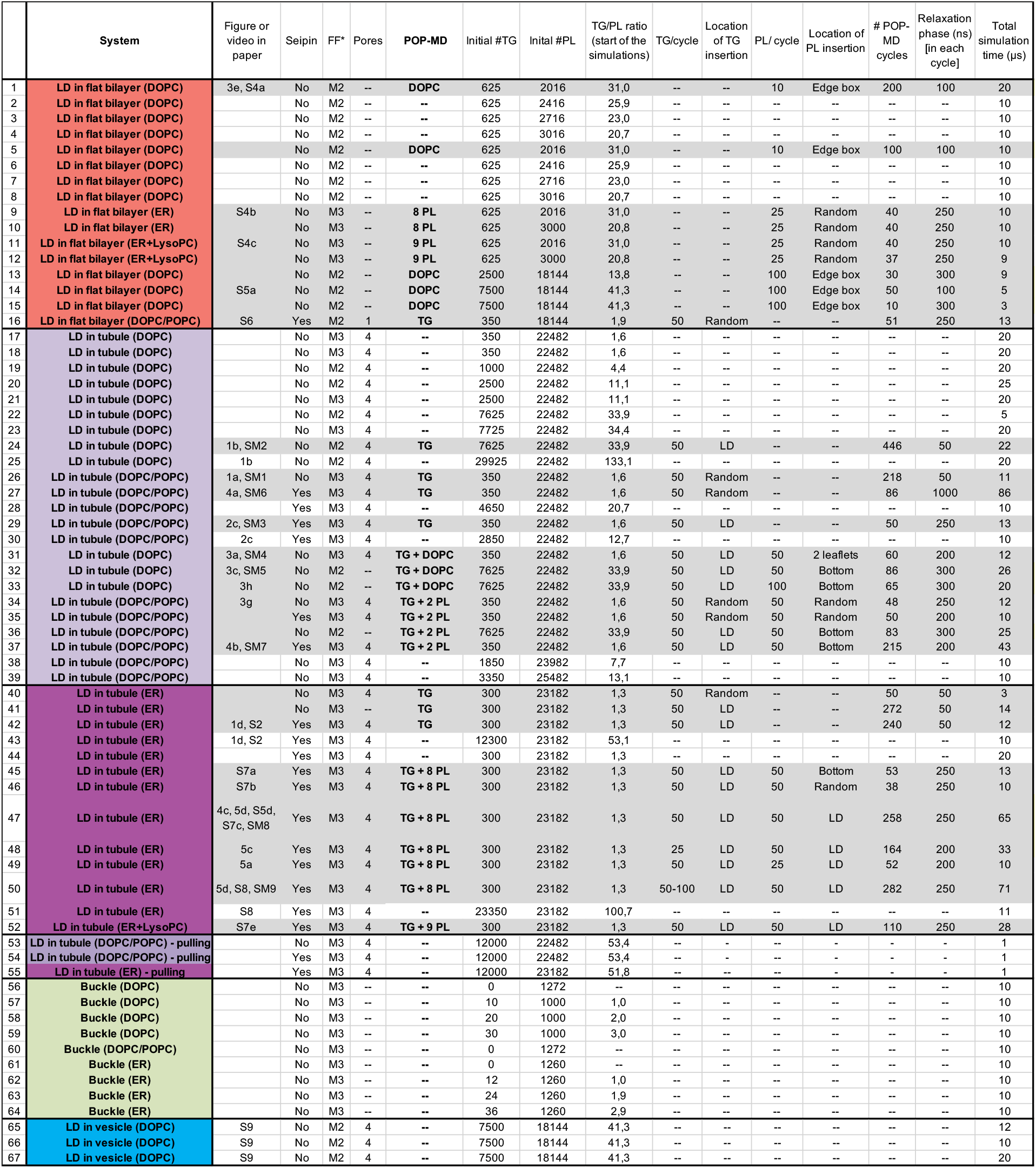
Summary of the main simulations described in this work. For each system, we report a simplified description, the Figure or Movie number (if present in the paper), the presence of seipin, the force field (M2 for Martini 2 and M3 for Martini 3), the number of hydrophilic pores (if present), the nature of the molecules inserted in the system by POP-MD, the number of TG and phospholipid (PL) molecules in the beginning of the simulation, the number of TG and PL molecules inserted by POP-MD and their location, the number of POP-MD cycles and their relaxation time, and the simulation time. In the POP-MD column, “2 PL” (or “8 PL”, “9PL”) indicates that 2 phospholipids (or 8, or 9, respectively) were inserted by POP-MD, in the same proportion as in the starting system. POP-MD simulations are shaded in gray.

**Figure 1.**
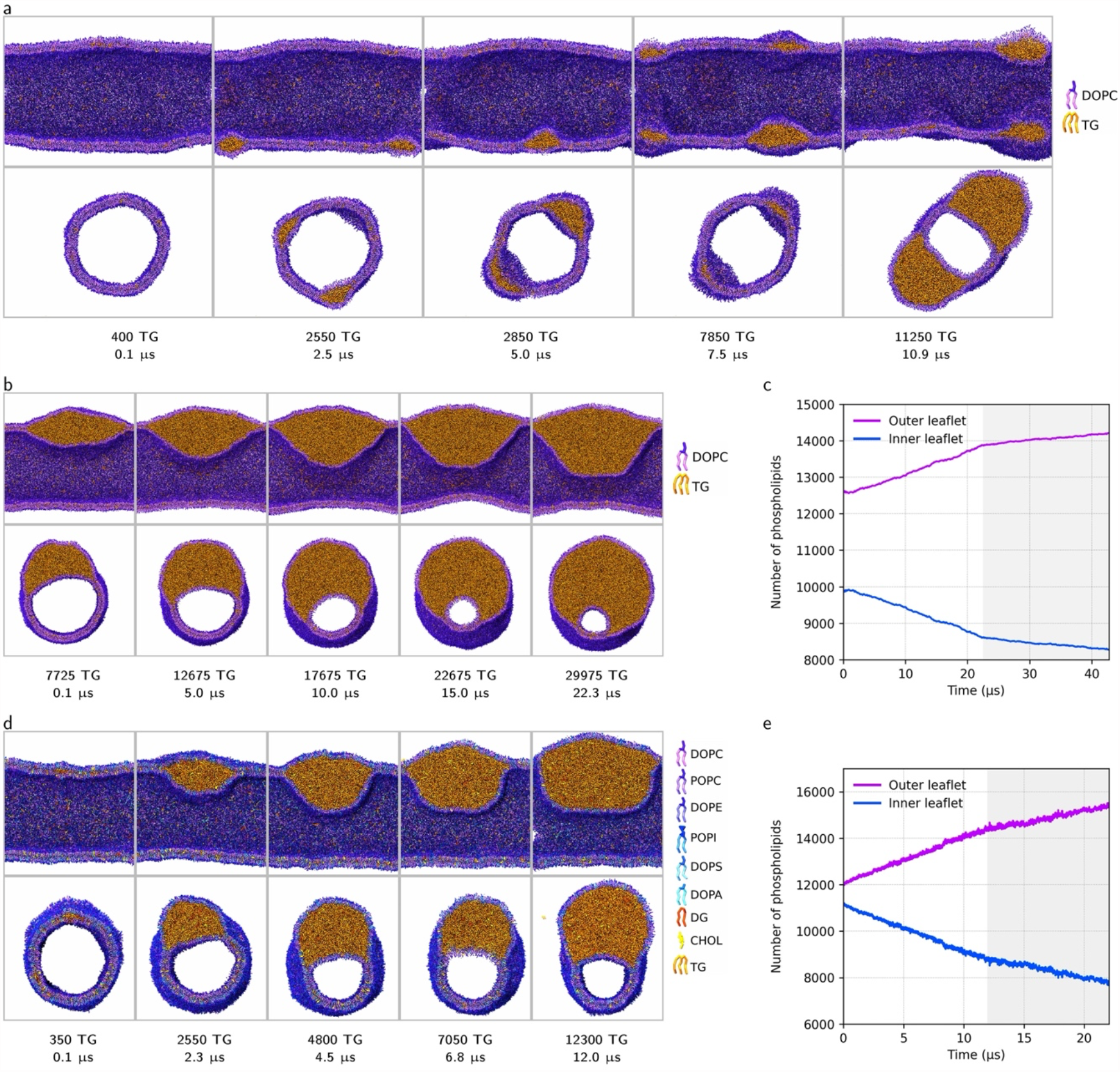
Growing LDs in tubular membranes by TG synthesis. (a) Snapshots from POP-MD simulations with a pure dioleoylphosphatidylcholine (DOPC) membrane tubule, with TG synthesis in random locations within the tubule; side view cut along the tubule and perpendicular to the tubule; the color code is in the legend on the right-hand side. Multiple nascent LDs are formed and do not coalesce on the simulation time scale. (b) Snapshots from POP-MD simulations with TG synthesis within or in the proximity of the nascent LD; in this case, only one nascent LD is formed. (c) Analysis of leaflet imbalance during POP-MD (0-22 μs) and subsequent equilibrium MD (22-42 μs, shaded area). (d) Snapshots from POP-MD performed on a tubule with ER-mimicking bilayer composition (see Methods), and (e) analysis of leaflet imbalance during POP-MD (0-12 μs) and the subsequent equilibrium MD (12-22 μs, shaded area).

Synthesis of TG in one specific region in the tubule induced nucleation of a single nascent LD, followed by growth, but no budding (Fig. 1b, Supplementary Movie 2). In fact, the LD protruded on both sides of the bilayer; the larger fraction of its volume protruded towards the lumen of the tubule, even if the number of phospholipids in the outer leaflet grew throughout the simulation (Fig. 1c). We repeated the POP-MD simulations using a more realistic lipid composition (Fig. 1d), mimicking the ER^24,25^, consisting of 8 different types of lipids (see Methods). Localized synthesis of TG promoted asymmetry between the leaflets in terms of number of lipids (Fig. 1e) and composition, with the outer leaflet progressively enriched in PC lipids and the inner leaflet in PE and PI lipids (Fig. S2). Despite the spontaneous asymmetry buildup, no budding was observed (Fig. 1d) and the LD protruded mostly towards the lumen of the tubule. Protrusion of the LD towards the lumen might appear surprising, as it rarely occurs in biological systems^5^, but can be understood based on the geometric constraints imposed by the tubule: protrusions *outwards* require an increase in the surface area of the outer leaflet, while protrusions *inwards* do not require any change in surface area. Emergence of cellular LDs towards the cytosol cannot be due to the tubular morphology: other asymmetric structures or processes must determine the direction of budding.

Besides growth of LD volume, different factors may be required for directional LD budding. First, changes in the relative surface area of each membrane leaflet, necessary for LD budding, may require large lipid reservoirs, available in the ER; according to theoretical work, area fluctuations enable spontaneous budding of small LDs (few tens of nm)^10,11,26^. Second, LD budding is observed in cells in the presence of seipin at the LD-bilayer junction^9^; seipin has a major role in LD biogenesis, and may be necessary for budding. Third, while surface tension is negligible in membrane tubules without TG, it is possible that TG synthesis increases the surface tension, which would prevent budding^27^. Last, while the ER is known to be one of the most symmetric membranes in cells^28^, local asymmetry may be required for LD budding. In the following, we will explore these possibilities, to understand which forces drive LD budding.

### Seipin scaffolds nascent LDs but is not sufficient to induce directional budding

Seipin is an oligomeric ER membrane protein critical for normal LD formation, as it defines LD nucleation sites^29^ and maintains LD-ER contact after budding^8,9,30^. The structure of seipin has been solved at high resolution by cryo-EM for human^31^, drosophila^32^, and yeast^33,34^ proteins, and consists of a highly conserved luminal domain, two transmembrane domains forming a ring of ∼15 nm in diameter, and one N-terminal cytosolic domain. It has been suggested that seipin favors LD nucleation by catalyzing TG aggregation and trapping TG within the perimeter defined by its TM helices^13-15,35^. It has also been proposed that seipin plays a major role in preventing abnormal phenotypes, with few super-sized LDs and numerous tiny ones, by decreasing the probability of LD growth by ripening^13,35^. To understand the role of seipin in the budding process, we simulated TG synthesis in the proximity of the protein, as observed in cells^30,36^, using the Drosophila protein (Fig. 2a-b, S3) and a membrane tubule with the ER-mimicking lipid composition. In the initial stages of growth, the nascent LD was scaffolded by seipin and acquired an asymmetric shape protruding towards the cytosolic compartment, not observed in the absence of the protein (Fig. 2c, 9 μs). However, once the size of the nascent LD exceeded the diameter of the protein, its transmembrane portions deformed (Fig. 2d), TG molecules leaked out of the scaffold, and the LD leaned towards the interior of the ER tubule. Leakage of TG was enabled by the high flexibility of seipin in the region connecting the luminal and transmembrane domains, consistent with experimental data^34^. Extension of the simulation without addition of TG produced a sort of budded shape, with the LD budded in the lumen (Fig. 2c, 23 μs, Supplementary Movie 3), not towards the cytosol.

**Figure 2.**
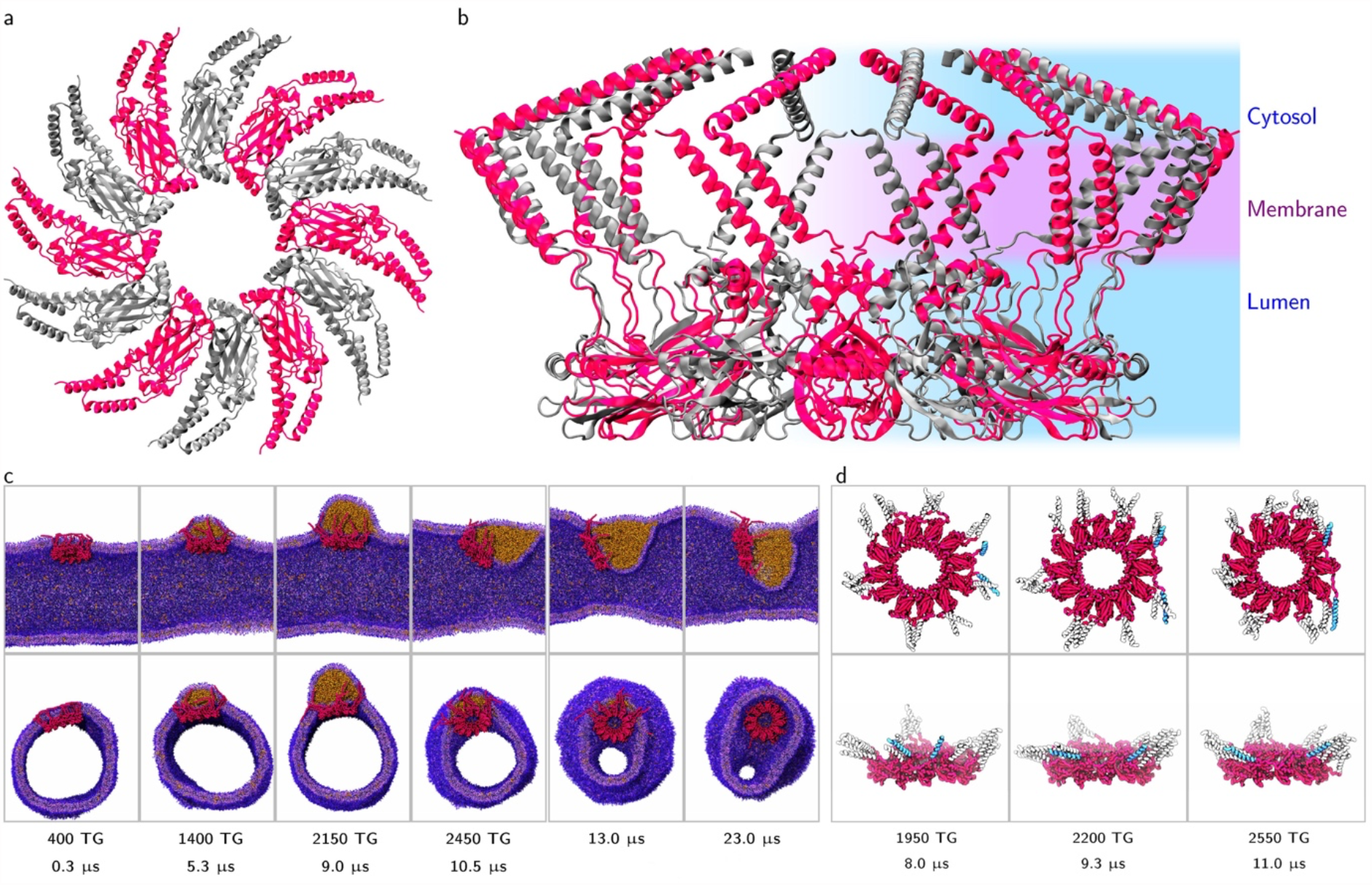
Growing a TG droplet in a tubular membrane in the presence of seipin. (a) Complete model of seipin, based on cryo-EM structure from Drosophila^32^, top view and (b) side view. Protomers are colored in magenta and grey to distinguish them. (c) Snapshots from LD growth simulations and subsequent equilibrium MD; seipin is in red, the color code for lipids is the same as in Fig. 1; the number of TG molecules inserted and the simulation time are indicated for each snapshot. (d) Seipin scaffold opening: the transmembrane helices (white or light blue) are initially equally spaced around the luminal domain (magenta); at time = 9.3 μs, two of them (in light blue) get separated, allowing TG to leak out of the seipin scaffold.

### Symmetric phospholipids synthesis induces budding towards the lumen

In ER tubular membranes, surface tension is negligible^20^, which makes LD budding favorable. In our simulations, surface tension was negligible in the absence of LDs (SI.4), as shown by the small contractive force along the tubule axis. In contrast, in the presence of LDs, the contractive force was significant and increased as the LD grew (SI.7), suggesting that surface tension increases during LD growth – which reduces the probability of budding^27^.

To guarantee low surface tension in the system during LD growth, we simulated simultaneous synthesis of TG (in the proximity of an existing LD) and phospholipids (in both bilayer leaflets). In this case, the force acting along the tubule remained approximately constant (Table S3), indicating that the symmetric insertion of phospholipids in the bilayer counteracts the effect of the TG droplet. Remarkably, symmetric phospholipid synthesis did induce a budding transition, with the LD volume entirely on one side of the bilayer, but the LD budded within the tubule (Fig. 3a, Supplementary Movie 4), as observed in the presence of seipin (Fig. 2c). Low surface tension is indeed crucial for LD budding, in agreement with theories^10,26^ and experiments^27^, but does not induce budding towards the cytosol.

**Figure 3.**
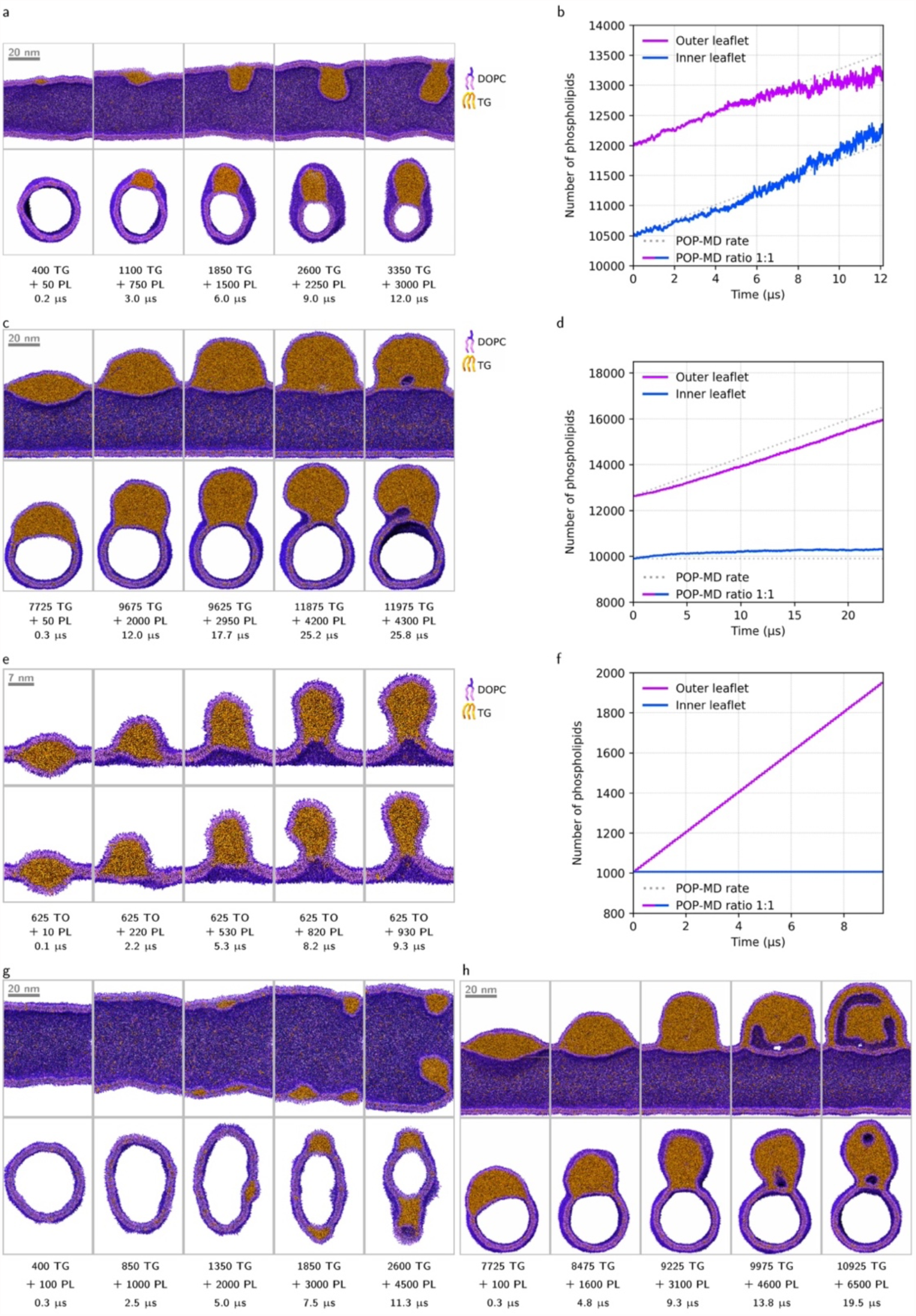
LD budding induced by synthesis of TG (close to the existing nascent LD) and phospholipids (PL), in the absence of seipin; total number of TG molecules, number of inserted PL molecules, and simulation time are indicated in each panel. (a) LD budding from a tubular membrane (22482 DOPC before POP-MD) by synthesis of DOPC lipids in both leaflets (TG:PL ratio 1:1). The *budding transition* generated structures qualitatively different from the ones observed without PL synthesis: the LD volume is entirely on one side of the cylindrical profile of the tubule. (b) Build-up of leaflet asymmetry during POP-MD with symmetric PL synthesis; dashed lines represent the theoretical number of lipids in the absence of flip-flops. (c) LD budding from a tubular membrane (22482 DOPC before POP-MD) by synthesis of TG (near the center of the tubule) and DOPC lipids only in the outer leaflet (TG:PL ratio 1:1), and (d) build-up of leaflet asymmetry. (e) LD budding in a small flat membrane (2016 DOPC lipids before POP-MD, ∼26 nm in lateral size, 625 TG molecules) by synthesis of DOPC only in the upper leaflet, and (f) build-up of leaflet asymmetry. (g) Synthesis of TG in random locations of a DOPC tubule gives multiple nucleated LDs. (h) Formation of water-filled defects at the LD-tubule connection (22482 PL before POP-MD, TG:PL ratio 1:2); defects protruding into the oil region are bounded by a phospholipid monolayer.

### Asymmetric phospholipid synthesis is necessary and sufficient for directional LD budding

According to experiments, theory, and simulations, leaflets asymmetry is necessary for LD directional budding^11,12^. In synthetic systems, this can be achieved by adding lipids (with or without positive intrinsic curvature^11,12^) to the outer leaflet of the bilayer, or by adding proteins that bind the outer LD monolayer^12^. In the ER, asymmetry can be generated by the synthesis of PC lipids in the outer leaflet by CCTα, an enzyme active when bound to (PC-deficient) LDs^37^. In our simulations, leaflet asymmetry developed spontaneously (due to lipid flip-flops, Fig. S2), but remained limited. To better mimic the biological process, we simulated simultaneous synthesis of TG and PC lipids, with PC added only to the outer leaflet of the tubule, to *actively* generate leaflet asymmetry while reducing surface tension. We finally did observe, for the first time, LD budding towards the cytosolic compartment (Fig. 3c, Supplementary Movie 5).

We wondered whether asymmetric synthesis of phospholipids is not only necessary but also sufficient for LD budding. To address this question, we repeated POP-MD simulations on very small systems with flat geometry, no TG synthesis, and simple composition (pure DOPC). Remarkably, we could reproduce LD budding on nascent LDs as small as 12 nm in diameter (i.e., less than 600 TG molecules), simply by adding DOPC lipids to one membrane leaflet (Fig. 3e). The tubular morphology, the large LD volume, the nature of the phospholipids, and the presence of seipin are *not* essential for budding (Fig. S4); only low surface tension and leaflet imbalance are necessary and sufficient for LD budding.

### Seipin yields a robust budding mechanism

The budding process simulated in the absence of seipin shows some unique features: the LD-tubule connection is large (Fig. 3c), only limited by the size of the tubule (∼35 nm), and cannot accommodate the presence of seipin; multiple nascent LDs nucleate when TG is synthesized in random locations (Fig. 3g), because LD nucleation is faster than LD coalescence; finally, large deformations are formed at the LD-tubule connection, with water-filled defects protruding into the core of the budded LD (Fig. 3h, S5). Are these features directly related to the role of seipin? To test this hypothesis, we repeated LD growth simulations in the presence of seipin, with asymmetric synthesis of phospholipids and TG synthesis in the proximity of seipin or away from it. A nascent LD formed rapidly within the seipin ring, and nucleation required lower TG concentration compared to tubules without seipin, as observed by others^13,14^. However, when TG was synthesized at random locations, multiple nucleation events were observed outside the seipin ring (Fig. 4a, S6, Supplementary Movie 6), suggesting that the favorable TG-seipin interactions may not be sufficient to avoid the formation of multiple LDs. When TG synthesis was localized in the proximity of seipin, budding was observed when the nascent LD became larger than the seipin ring (Fig. 4b, Supplementary Movie 7). Seipin remained at the LD-ER contact site, constraining its size (15 nm) and thereby endowing the budded LD with a lightbulb shape. The constraint on neck size originates from the protein amino acid sequence: hydrophobic transmembrane helices are linked to the luminal domain by short loops and are bounded by charged residues or hydrophilic loops at both ends; the helices cannot get far from the luminal domain and crossing the membrane would imply a prohibitive (electrostatic) energy barrier, which prevents bilayer unzipping, in agreement with recent simulations^15^. Remarkably, no water-filled membrane defects were formed in the proximity of the LD neck. Instead, the cylindrical shape squeezed to avoid the formation of high-curvature regions in the proximity of LD neck (Fig. 4b). Overall, seipin had a clear effect on the mechanism of budding, yielding an approximately spherical budded LDs with a narrow, stable, and defect-free neck.

**Figure 4.**
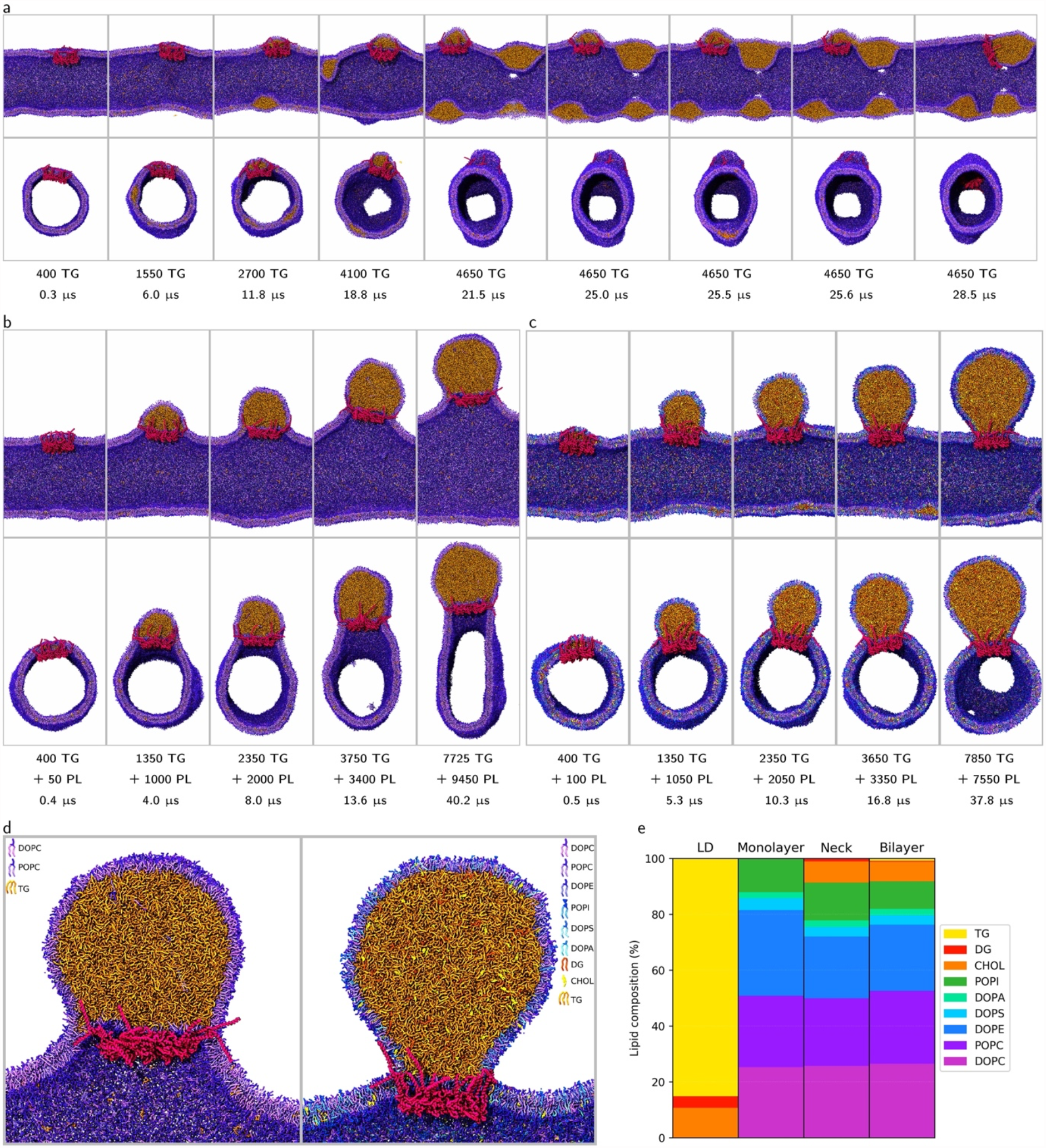
Growing a TG droplet in a tubular membrane in the presence of seipin by synthesis of TG and asymmetric synthesis of PL (only in the outer leaflet), in tubules made of PC lipids only (DOPC:POPC 1:1), with TG synthesis (a) in random locations and (b) in the proximity of seipin; (c) in tubules with the complex ER lipid composition (TG synthesis close to seipin, PL synthesis only in the outer leaflet, TG:PL ratio 1:1); the total number of TG, the number of inserted PL molecules, and the corresponding simulation time are indicated in each panel. (d) Close-up of the LD-tubule connection (i.e., the LD neck) with pure PC tubules (left panel) and the ER lipid mixture (right panel); the number of TG molecules is similar (7725 vs 7850) but the ER mixture also contains cholesterol and DG, which distribute in part to the oil region of the LD, increasing its volume. (e) Average lipid composition of the different regions in the system in systems with the ER lipid composition, in equilibrium simulations following LD budding. Cholesterol, DG, and POPI are enriched in the LD neck compared to the bulk bilayer region; the oil region of the LD contains a significant fraction of cholesterol and DG.

### The complex ER composition stabilizes LD-ER contact sites

We could simulate LD budding using a very simple lipid composition: only one type of neutral lipid and PC phospholipids (Fig. 3, 4b). So, what is the role of the complex ER composition in the budding process? To address this question, we grew LDs in tubular membranes with ER-mimicking lipid composition^18,19^. The rich ER composition had no major effect on LD nucleation and growth, but budding produced structures with high-curvature LD necks: the tubule remained cylindrical throughout the budding process and the LD had a lightbulb shape (Fig. 4c, Supplementary Movie 8), in contrast with pure PC membranes (Fig. 4d). Two factors concur to this effect: the lower bending rigidity provided by the rich ER composition (∼11 k_B_T, compared to ∼18 k_B_T for pure DOPC), and the inhomogeneous lipid distribution around the protein (Fig. 4e), enriched in cholesterol, dioleoylglycerol (DG), and phosphatidylinositol (PI). Interaction of seipin with PI and PA lipids has been observed experimentally^31^, and is probably related to the positive electrostatic charge on residues adjacent to the transmembrane region. Cholesterol and DG, having smaller polar heads, fit well in the highly curved LD neck region, stabilizing it.

### Regulation of phospholipid synthesis

In cells, PC synthesis is regulated through a feedback loop by CCTα, and is tuned based on the surface tension in the ER membrane^37^. Regulation of TG synthesis in cells is crucial for LD production, but the reason for tight regulation of PC synthesis is less clear. We repeated LD growth simulations using different TG:PL ratios. High TG:PL ratio (2:1) resulted in failure to bud towards the cytosol, even if leaflet asymmetry could grow beyond the content supplied by POP-MD (Fig. 5a-b). With a low TG:PL ratio (1:2), tether-like bilayer deformations appeared in the ER tubule, away from the LD monolayer and LD-tubule contact site (Fig. 5c). Similar defects also formed with TG:PL 1:1 ratio, at later stages of LD growth (Fig. S5), and were completely avoided in simulations with variable synthesis rate, starting from TG:PL 1:1 and then increasing to TG:PL 2:1 after the budding transition (Fig. 5d, Supplementary Movie 9). The need to regulate PL synthesis (by increasing TG:PL ratio during LD growth) is explained by a simple geometric argument: the number of TG molecules depends on the volume of the LD, which grows as the cube of the (linear) size of the LD, while the number of PL depends on the surface area of the LD, which grows as the square of the LD size. As LDs grow, the cell needs to synthesize more TG than PL to maintain constant surface coverage of the LD. While the actual rates of addition of TG and PL used in our simulations are inconsequential to biological systems, the high sensitivity of the budding mechanism to PL synthesis rates explains why precise regulation of PL synthesis is crucial for LD biogenesis and for the stability of LD-ER contact sites.

**Figure 5.**
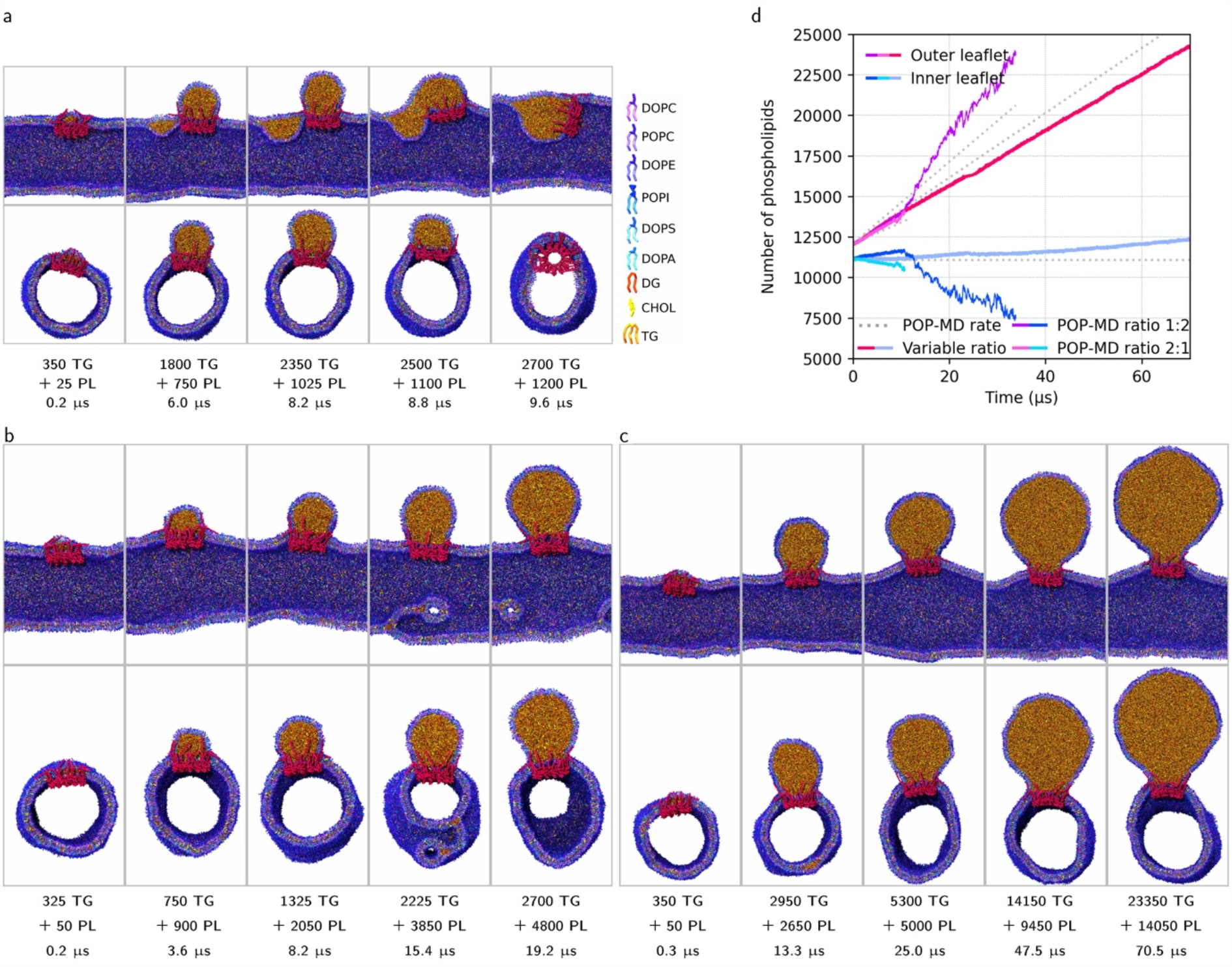
Growing a TG droplet in a tubular membrane with a complex ER mixture by adding both TG and 8 different PL types, in the presence of seipin. Snapshots from POP-MD simulations with different TG:PL synthesis ratios: (a) TG:PL 2:1 (b) TG:PL 1:2, and (c) variable ratio (starting with 1:1 and continuing with 2:1). (d) Build-up of leaflet asymmetry; the dotted lines represent the amount of PL synthesized by POP-MD in each leaflet; deviations from the dotted lines are due to spontaneous flip-flop (due to hydrophilic pores).

Finally, we asked the question on the localization of PL synthesis and its effect on budding mechanism and LD morphology. Synthesis of phospholipids close to seipin, away from seipin, or in random locations (in the outer leaflet of the tubule) resulted in all cases in a single nucleated LD, that budded towards the cytosol; the lightbulb shape of the LDs was virtually identical, independent of the localization of PL synthesis (Fig. S7). Localization of PL synthesis is irrelevant for the budding mechanism, in striking contrast with the localization of TG synthesis, strongly affecting the number of nucleated LDs.

## Discussion

The mechanism of LD biogenesis is difficult to explore via current structural biology techniques, because of the fluid nature of biological membranes and LDs, as well as limitations in spatial and temporal resolution. Theory and simulations provided important contributions to our understanding of LD biogenesis^7,10,11,13-15,23,38^, but the current picture is far from complete. LD nucleation and phase separation were observed in simulations before^7,13,22,23,38^, and occur on fast time scales (below the microsecond); in contrast, the budding process has never been observed so far, neither in simulations nor experimentally. Due to this gap of knowledge, questions on the role of different proteins and lipids and even on the driving forces for LD budding remained open so far.

Here we addressed open questions on the driving forces and the mechanism of LD budding, as well as on the regulation of lipid synthesis and on the role of seipin. To this end, we developed a novel computational method, coined POP-MD, that allows molecular-level simulations of LD growth by emulating lipid synthesis. POP-MD enables modeling the entire mechanism of LD biogenesis, from nucleation in ER tubules to growth and budding. Simulations of LD growth allow, first and foremost, to address the central question on the driving forces for LD budding. The current view is that synthesis of neutral lipids, necessary for LD nucleation and growth, is sufficient to also drive the budding process^10,26,39^; this is because the presence of a nascent LD causes an increase in the surface tension in the LD monolayer region^27^ (while the bilayer remains at very low tension^20,40^), and LD monolayer surface tension drives budding^27^. However, experimental validation is limited. Our simulations show, instead, that nascent LDs *spontaneously bud towards the lumen* in tubular membranes, when surface tension in the tubular bilayer is sufficiently low; this is because budding *towards the cytosol* requires a significant increase in the surface area of the outer leaflet of the ER membrane, while budding *towards the lumen* does not. Therefore, asymmetric synthesis of phospholipids is necessary for budding towards the cytosol. More precisely, low surface tension in the tubular bilayer and leaflet asymmetry are necessary and sufficient for directional budding. A corollary to this result is the absence of a well-defined size threshold for the budding transition: large LDs (over 50 nm in size) would not bud in symmetric membranes, while much smaller ones (∼12 nm) bud spontaneously when leaflet asymmetry is sufficient. The apparent discrepancy between theoretical and simulation results stems from different assumptions: in continuum theories, the bilayer is assumed to be of infinite size, hence it acts as a phospholipid reservoir and allows *local* leaflet asymmetry, via surface area fluctuations; in simulations, instead, reservoirs are small, which makes it necessary to actively generate leaflets asymmetry. Is the assumption of a large reservoir realistic for the ER tubular network, or rather asymmetric synthesis explains directional budding? We contend that simulations offer a more realistic picture, because in cells, in the absence of phospholipid synthesis, LDs remain trapped in the ER membrane^41^; this event is associated with conditions that alter phospholipid synthesis or degradation^41^. Differences in composition between LD monolayers and ER bilayers, observed both experimentally^42^ and in our simulations (Fig. 4e, S8), are also consistent with asymmetric PL synthesis during budding.

The view of leaflet asymmetry emerging from our simulations is consistent with experimental data, indicating that membrane asymmetry determines the direction of LD budding from the ER^12^. However, simulations show a much more detailed picture of the budding mechanism: they predict the existence of a *budding transition* as the key step in the process. Such prediction is amenable to experimental validation, although trapping the different stages of budding is challenging for current microscopy techniques. The shape and diameter of the LD neck depend on the presence of seipin: only in the presence of seipin we observed the lightbulb shape previously hypothesized^10,26,39^. The geometric features of the LD-tubule connection spontaneously generated in simulations match very well cryo-electron tomograms of LD-nuclear envelop contact sites^35^.

Since simulations provide information on driving forces and morphologies, it is tempting to also consider the kinetics of budding. However, the budding kinetics in simulations is determined by the (user-defined) PL synthesis rate, chosen to be extremely high (between 0.1 and 1 lipid per ns) to reduce the computational cost; even considering that many proteins concur to lipid synthesis along an ER tubule, the synthesis rates used in our simulations exceed the turnover of lipid-synthesizing enzymes by several orders of magnitude. While lipid synthesis rates were user-defined, the response of nascent LD systems to the buildup of leaflet asymmetry was unbiased and occurring on fast time scales, below the microsecond, suggesting that, *in vivo*, the rate of the initial steps in LD budding depends not only on triglyceride synthesis rates, but also on phospholipid synthesis rates.

The detailed picture of the LD budding mechanism obtained in different conditions allows us to address a number of biological questions and make testable predictions: on the role of seipin in the budding process, the role of ER morphology and lipid composition, and the regulation of TG and phospholipid synthesis.

- In cells, LDs generally form in tubular regions of the ER^17^, but the tubular morphology favors budding towards the ER lumen (Fig. 3a); if phospholipid synthesis in the outer leaflet of the ER is impaired, LDs should bud towards the lumen, not the cytosol.
- LDs show similar budding propensity in flat and tubular membranes (Fig. 3e, S4); budding from flat regions of the ER or the nuclear envelope is predicted to follow the same mechanism as in the tubular ER. We predict that preferential formation of LDs in tubular regions is related not to different budding propensities, but to different nucleation propensities and/or distribution of proteins.
- Seipin scaffolds the LD neck, as previously proposed based on a cryo-EM structure^34^ and equilibrium simulations^15^; but it cannot promote budding towards the cytosol when leaflet asymmetry is insufficient (Fig. 2c).
- Seipin favors TG condensation^13,14^ but, *per se*, cannot prevent the formation of multiple LDs when TG is synthesized in random locations (Fig. 4a), due to fast LD nucleation and slow LD coalescence. LD biogenesis in cells is orders of magnitude slower than in simulations, but the length of ER tubules is orders of magnitude greater, and larger nascent LDs will diffuse more slowly; we predict that coalescence will be slow also in cells. Also, we propose that seipin reduces the number of nucleated LDs by localizing TG synthesis in its proximity, possibly by recruiting TG-synthesizing enzymes. Recruitment of TG synthesis enzymes in the proximity of seipin has been observed in yeast^30^. Nucleation of multiple LDs matches experimental observations in seipin-knockout yeast and human cells, resulting in aberrant phenotypes with few giant LDs and numerous tiny LDs^8,29,43^.
- Seipin prevents the formation of defects in the proximity of the LD neck. We speculate that monolayer-bounded defects may act as diffusion barriers for proteins, therefore altering the protein composition of the LD surface. However, we notice that both monolayer bounded defects in the LD neck (Fig. 3h, S5a) and bilayer bounded defects in the tubules (Fig. S5d) are the result of excessive surface density of the outer leaflet; a “normal phenotype” (i.e., absence of deformations) can be rescued by tuning the TG:PL ratio during the simulations; this is analogous to the way cells operate, by regulating PC synthesis based on the surface density (i.e., the surface tension) in the ER outer leaflet^37^. Therefore, both types of defects are unlikely to form in cells, as phospholipid synthesis stops when surface density is high.
- The rich ER lipid composition favors budding and reduces deformations of tubular membranes due to low bending rigidity and lipid sorting in the proximity of the seipin ring (Fig. 4e) and the LD monolayer (Fig. S8). Our simulation results do not imply that synthesis of *all* phospholipids is important for the budding mechanism: in cells, synthesis of a single type of phospholipid (e.g., PC) would generate asymmetry in surface density without significantly altering the overall ER composition, because the extent of the ER membrane is far superior to the surface area of the LD. Differences in the composition of the LD monolayer compared to the ER tubule are compatible with available mass spectrometry data^42^.
- Regulation of PL synthesis and TG:PL ratio is predicted to be crucial for the budding mechanism: it determines the direction of budding, the onset of the budding transition, and the appearance of deformations and defects. Localization of TG synthesis close to seipin is necessary to reduce the number of nucleated LDs (Fig. 4a), and is consistent with recent findings on the interaction of DGAT2 (an enzyme synthesizing TG) with LDs^44^. Localization of phospholipid synthesis, instead, has no effect on the budding mechanism (Fig. S7), and does not require spatial coupling with seipin or LDs. Remarkably, this is also compatible with localization of CCTα in the nuclear envelope (NE), observed in experiments^45^; as the NE is continuous with the ER, PC synthesis in the NE would yield the same LD budding mechanism. Our simulations explain why localization of TG synthesis is important while localization of PL synthesis is not: coalescence of multiple nucleated LDs requires diffusion of individual TG molecules (growth by Ostwald ripening^39^) or TG droplets (growth by fusion); individual TG molecules diffuse a few μm^2^/s, TG lenses much less, while changes in surface density (and surface tension) propagate very fast (meters per second, Fig. S10).

Simulations of LD growth reported here were enabled by the POP-MD workflow, emulating lipid synthesis. Depending on the synthesis rate, the simulated systems may or may not reach equilibrium during the relaxation phase. The lack of equilibrium is problematic for the calculation of thermodynamic properties (e.g., surface tension) of the simulated systems, which is a limitation of the methodology. On the other hand, the morphological features and transformations of nascent LDs proved to be robust and reproducible, as they depend mostly on surface densities, which propagate fast on simulation time scales. We anticipate that our methodology will enable simulations of even more complex transformations of membrane systems, including the biogenesis of other organelles, the formation of viral envelopes, and bacterial division.

## Author contributions

LM designed the research. VN wrote the POP-MD simulation protocol. VN and JC performed and analyzed most simulations. DS wrote and tested customized versions of Suave. VN prepared most figures. All authors analyzed and discussed the results. LM wrote the manuscript. All authors reviewed and edited the manuscript.

## Acknowledgement

The authors thank Samuli Ollila, Lionel Foret, and Abdou Rachid Thiam for discussions on the physics of lipid droplets, Nika Zarubina and Juliette Martin for support in building the model of seipin. All calculations were performed at French supercomputing centers (CINES, IDRIS, and TGCC) supported by *Grand Equipement National de Calcul Intensif* (GENCI, grant number A0080710138, A0100710138, A0120710138). We acknowledge CC-IN2P3 (https://cc.in2p3.fr) for computing services (data storage and backup). LM acknowledges funding by the *Institut National de la Santé et de la Recherche Médicale* (INSERM) and ANR (grant ANR17-CE11-0003-01, ANR21-CE11-0032-01).

## Methods

All simulations used the Martini coarse-grained force field, either version 2^46,47^ or version 3^48^, as specified in Table 1, and were carried out with the Gromacs^49^ software (version 2018 or 2019). Calculations of local stress tensor relied on the Gromacs-LS package^50^, and were carried out only on equilibrium simulations. In the following, we first describe the simulation setup for the different systems studied here, while simulation parameters are provided in the Supporting Information.

### Setup of nascent LDs in flat bilayer systems

We generated flat periodic lipid bilayer systems using the Insane software^51^ or the MAD web server^52^. The dimensions of the bilayer ranged from 27×27 nm (2016 phospholipids) to 78×78 nm (18144 phospholipids); a complete list of the simulated systems is reported in Table 1. To generate nascent LDs, we initially built tri-layer systems, with a layer of triglycerides (TG) sandwiched between two layers of phospholipids. Pure DOPC was used for the bilayer, while pure triolein was used as the triglyceride (oil phase). Additional simulations of flat bilayers with nascent LDs (SI.8, Fig. S4) used a more complex ER-mimicking composition, containing 25% dioleoyl-phosphatidylcholine (DOPC), 25% palmitoyl-oleoyl-phosphatidylcholine (POPC), 23% dioleoyl-phosphatidylethanolamine (DOPE), 3% dioleoyl-phosphatidylserine (DOPS), 2% dioleoyl-phosphatidic acid (DOPA), 10% palmitoyl-oleoyl-phosphatidylinositol (POPI), 10% cholesterol (CHOL), and 2% of dioleoyl-glycerol (DG).

### Setup of vesicular and tubular systems with hydrophilic pores

A vesicle with an embedded LD (18144 DOPC and 7500 TG) was generated from a flat bilayer by removing membrane periodicity (by adding water around the membrane); line tension rapidly transformed the square membrane in a circular one, that bent out of plane to reduce the perimeter of the open edge and eventually closed, forming a vesicle.

Tubular membranes were then obtained from the vesicle by generating large water pores on opposite sides of the vesicle (along the *x* axis) and then connecting the periodic images by adding lipids with POP-MD (see below). Pure DOPC tubules contained 22482 lipids, had an external diameter of 35 nm, length of 75 nm, and were periodic in the direction of their main axis (*x* axis). We also built tubules with DOPC:POPC 1:1 mixtures and with a more complex composition, mimicking the ER composition^24,25^ (see above), by replacing DOPC and/or by adding other lipids with POP-MD (see below). The complex systems contained 5620 DOPC, 5620 POPC, 5170 DOPE, 676 DOPS, 450 DOPA, 2248 POPI, 2248 CHOL, and 450 DG molecules, for a total number of 23182 lipids. For all tubular membranes, the size of the system was approximately 75×50×70 nm (or larger in the *z* dimension in the case of steered MD simulations, see SI.1), with about 2.2 million particles before POP-MD runs (and up to 3 million particles at the end of POP-MD runs, after LD growth). The length of the tubule was always fixed in the *x* direction (tubule axis) and the *y* direction, while pressure coupling was applied in the *z* direction.

Tubular membranes contained four stable circular pores, generated using flat-bottomed potentials, as implemented in Gromacs^49^; flat bottomed potentials acted only on acyl chains, restraining them out of cylindrical regions in the membrane, and induced the formation of hydrophilic pores (i.e., with lipid head groups in contact with water). Hydrophilic pores make lipid flip-flop barrierless, allow dissipation of asymmetric stresses and differences in pressure in/out of the tubule (water can freely flow in/out), as well as changes in tubule diameter and surface area (Fig. S1). This setup leads to negligible surface tension, in the absence of nascent LDs (SI.4), effectively mimicking biological systems^20^.

### Models of seipin

We built a model of seipin dodecamer based on the Drosophila monomer structure (pdb code 6MLU^32^), adding the transmembrane (TM) and the N-terminal helices based on secondary structure prediction algorithms and AlphaFold2^53^. The putative N-terminal helix has been predicted but not resolved in any of the cryo-EM structures. The two transmembrane domains, instead, are resolved in the cryo-EM structure of yeast seipin^33^. Comparison with the structure of yeast seipin shows high similarity of both the luminal domain and the TM segments (Fig. S3). In the model, secondary and tertiary structure of the protein were maintained using elastic networks^54^ in the luminal domain, while no elastic network was required to fix the distance among 24 TM helices, because the loops connecting them to the luminal domain are short (11 amino acid residues).

### POP-MD simulations

Current MD software does not allow, to the best of our knowledge, changes in system composition “on-the-fly”. However, a simple way to achieve this is via a “stop-and-go” procedure: a simulation is performed in the desired thermodynamic ensemble, then stopped after a defined time; particles are inserted into the system, and the simulation is resumed. We devised an iterative procedure, coined POP-MD, which can insert any (small) molecule in user-defined regions of the system, emulating the synthesis of specific molecules at specific locations in the system. The POP-MD protocol consists of a non-equilibrium growth phase, in which lipids (or any other molecules) pop into the system, and a relaxation phase, in which the system drifts towards a new equilibrium (see SI). The duration of the growth and relaxation phase is user defined. After relaxation, the cycle is repeated, until the system size reaches a target value. If the relaxation phase is long enough, the properties of the system (energy, pressure, fraction of free TG, lipid distribution, etc.) may reach equilibrium. In principle, addition of lipids could be achieved at the same rate as determined by lipid synthesis in cells. However, considering a turnover rate of 10^4^ s^-1^ for a typical enzyme, this is much too slow for present-day supercomputers. On the other hand, very short relaxation runs between subsequent POP-MD steps pose problems in terms of computational stability, as systems need to adjust their volume after each addition of lipids, lipids and water need to redistribute by diffusion to dissipate internal stresses in the membranes and possible differences in pressure in/out of the tubule. Therefore, a compromise needs to be found between realistic growth rates, computational efficiency, numerical stability, and relaxation of stresses. We found that addition rates between 50 and 1000 lipids per microsecond result in numerically stable simulations (see Table 1). However, equilibrium is generally not reached in our simulations, in analogy with the biological systems we aim to understand, that are pushed out of equilibrium by lipid synthesis.

We carried out POP-MD simulations starting from either flat bilayer systems or from tubular membranes, with and without seipin. Hydrophilic pores were maintained open using flat-bottomed potentials. POP-MD works with and without hydrophilic pores, independently of system composition and force field.

Table 1 contains a list of the simulations discussed in the manuscript, the corresponding figures illustrating them, and the main features of the simulated systems (duration, POP-MD synthesis rate, system size, presence of seipin, hydrophilic pores, etc.). POP-MD simulations are shaded in grey, flat bilayer systems in red, tubular membranes in purple, vesicles in blue.

The total sampling, including all flat bilayer and tubular systems, was over 1 millisecond, and the overall computational cost was over 120 million CPU hours.

## Data availability

The starting structures, topologies, and molecular dynamics parameters files for 9 of the systems used in the current study are available on Zenodo: 10.5281/zenodo.8199687.

