## Supplementary Information for "Birth of an organelle: molecular mechanism of lipid droplet biogenesis"

### Supporting Information

##### 1. Simulations parameters

Standard Martini simulation parameters were used in all simulations; the Verlet neighborlist scheme[1] was applied, with a cutoff of 1.1 nm for non-bonded interactions and the Verlet cutoff scheme for potential shift[2]. Reaction field[3] was used in all simulations, with $\varepsilon_r$=15.
All MD simulations were carried out with the Gromacs[4] software (version 2018 or 2019) with the leap-frog integrator[5], except for simulations of buckled membranes, for which we used the Langevin integrator (*sd* in Gromacs). In all cases, the time step was 20 fs. In MD simulations, the temperature was kept constant with the Bussi-Donadio-Parrinello thermostat[6] (*v-rescale* in Gromacs, with time constant of 1 ps). In POP-MD simulations, the pressure was kept constant with the Berendsen barostat[7], with semi-isotropic pressure coupling (time constant of 12 ps, compressibility of 0 in the *xy* plane and $4 \cdot 10^{-5}$ bar$^{-1}$ in the *z* direction). In equilibrium simulations, the pressure was kept constant with the Parrinello-Rahman barostat[8] (with the same compressibility and time constants).

As detailed in the Methods section of the main text and in Table 1, most simulations contained stable pores in the bilayer membrane. The pores were generated using flat-bottomed potentials, as implemented in Gromacs[4], restraining lipid acyl chains out of cylindrical regions in the membrane. Flat-bottomed potentials used a force constant of 5000 KJ mol$^{-1}$ nm$^{-2}$ and a diameter of 5 nm. When LDs were present in the membrane, the pores were generated away from the LD.

POP-MD is a new protocol based on existing simulation codes and customized workflows, simulating the synthesis of lipids (and, in principle, any small molecules) by inserting them in the system at user-defined locations (for instance, within a given distance from the center or mass of a group of atoms, or within a given distance from a specific set of coordinates, or in a random location within the bilayer membrane, or within one membrane leaflet, etc.) and user-defined rates.
The protocol features five steps: 1, section of molecules to be inserted and their approximate localization; 2, insertion of new molecules with *gmx insert-molecules* (part of the Gromacs package); 3, slow growth MD simulation with position restraints (implemented in Gromacs) on newly inserted molecules (optional); 4, energy minimization; 5, relaxation phase (standard

MD). After relaxation, the cycle is repeated, until the system size reaches a target value. The number of lipids inserted at each cycle and the duration of the relaxation phase are user-defined.

In slow growth simulations, interactions (both Van der Waals and Coulomb) of the newly inserted molecules with all other molecules in the system were switched on progressively, by growing linearly the coupling parameter, $\lambda$, from an initial state $\lambda=0$, incremented by 0.0002 per time step (20 fs), so full coupling was achieved in 100 ps. The soft-core alpha and sigma parameters were set to 1.3 and 0.47 nm, respectively, while the soft-core power was set to 1.

Steered MD simulations were carried out on systems with LDs embedded in a vesicle or in a tubule, pulling the LDs away from the vesicle or tubule. Center of mass (COM) pulling was used, allowing to apply a force to the COM of a group of particles. A harmonic potential with fixed position was applied to the COM of the phospholipids (force constant of 2200 KJ $mol^{-1}$ $nm^{-2}$) to keep the vesicle or tubule at an approximately fixed position in the simulation box. A second harmonic potential was applied to the COM of TG, while increasing the distance between the COM of TG and the COM of the PL at different speed (0.5 nm/µs or faster; force constant of 2200 KJ $mol^{-1}$ $nm^{-2}$), to pull the LD away from the vesicle or tubule. The simulations were run for 1 µs or longer (Table 1). Additional simulations were started from the final frame of the steered MD run, keeping a fixed distance between the COM of the phospholipids and the COM of TG (force constant of 2200 KJ $mol^{-1}$ $nm^{-2}$). Finally, the force was released. Hydrophilic pores were kept open using flat-bottomed potentials as described above.

Analysis of the number of oil molecules in the nascent LDs was carried out with the g_aggregate software[9], while the number of lipids in each leaflet was calculated with the SuAVE software[10,11]. In SuAVE, the fitting process established in radial basis functions was used with the same parameters developed for closed surfaces[11]. A roughness parameter of 1 and a point resolution of 100 were used to define the grid points that make up the fitting surfaces. Then, a classification process was performed based on the distance between the lipids and the bidimensional fitted surfaces. A 2.5 nm cutoff was applied to distinguish the lipids from the different leaflets.

#### 2. Calculations of bending modulus from buckled membranes

We used the buckling method by Deserno[12] to calculate the bending modulus for three different lipid bilayer compositions: pure DOPC, 1:1 DOPC:POPC mixture, and the complex mixture mimicking the ER composition. In all cases we used identical methodology and parameters validated in our previous study[13]. Briefly, the starting bilayer structure was at least 32 nm x 8 nm x 20 nm in *x*, *y*, and *z*, in order to avoid finite size effects. The bilayer was then compressed by 30%, giving the system a corresponding strain ($\gamma$) of 0.3. Two production simulations for each system were then performed; the first, starting from the uncompressed, equilibrated bilayer, with no pressure coupling in *y*, allowing the bilayer to fluctuate only in *x*; the second being the compressed bilayer, with no pressure coupling in *x* or *y*, fixing the bilayer in its buckled state (Fig. S3). The uncompressed simulation provides the average *x* dimension length ($L$), and the buckled simulation allows for the calculation of the force exerted by the bilayer in the x-dimension of the buckle ($F_x$). From this, the bending modulus $k_c$ is calculated from the equation[12]:

$$F_x(\gamma) = 4\pi^2 k_c \frac{L_y}{L^2} \sum_{i=0}^{\infty} b_i \gamma^i$$

where the coefficients $b_i$ are reported in the original reference (ref. 12).
Composition of the ER mixture: 25% DOPC, 25% POPC, 23% DOPE, 3% DOPS, 2% DOPA, 10% POPI, 10% cholesterol, and 2% of DAG.

**Table S1.** Bending modulus obtained from simulations of buckled membranes.

| **Composition** | **$k_c$ ($k_BT$)** |
|---|---|
| pure DOPC | 18.5 ± 1.2 |
| DOPC/POPC 1:1 | 16.5 ± 2 |
| ER mixture | 11.2 ± 1.2 |
| DOPC + 1% TG | 18.0 ± 1.3 |
| DOPC + 2% TG | 16.5 ± 1.6 |
| DOPC + 3% TG | 15.4± 0.8 |

#### 3. Hydrophilic pores allow changes in tube diameter and surface area of each leaflet

We set up simulations of bilayer membrane tubules with hydrophilic pores, that were kept open by an external potential as detailed above. The presence of stable hydrophilic pores in membrane tubules allows barrierless flip-flop, which in turn dissipates asymmetric stresses and differences in pressure in/out of the tubule, and allows and changes in tubule diameter by pulling on the tubule with external forces. We confirmed this by generating tubules of different diameter simply by extending the box X dimension (along the tubule symmetry axis), Fig. S1. This setup also allows changes in the relative surface area of both leaflets, necessary for budding.

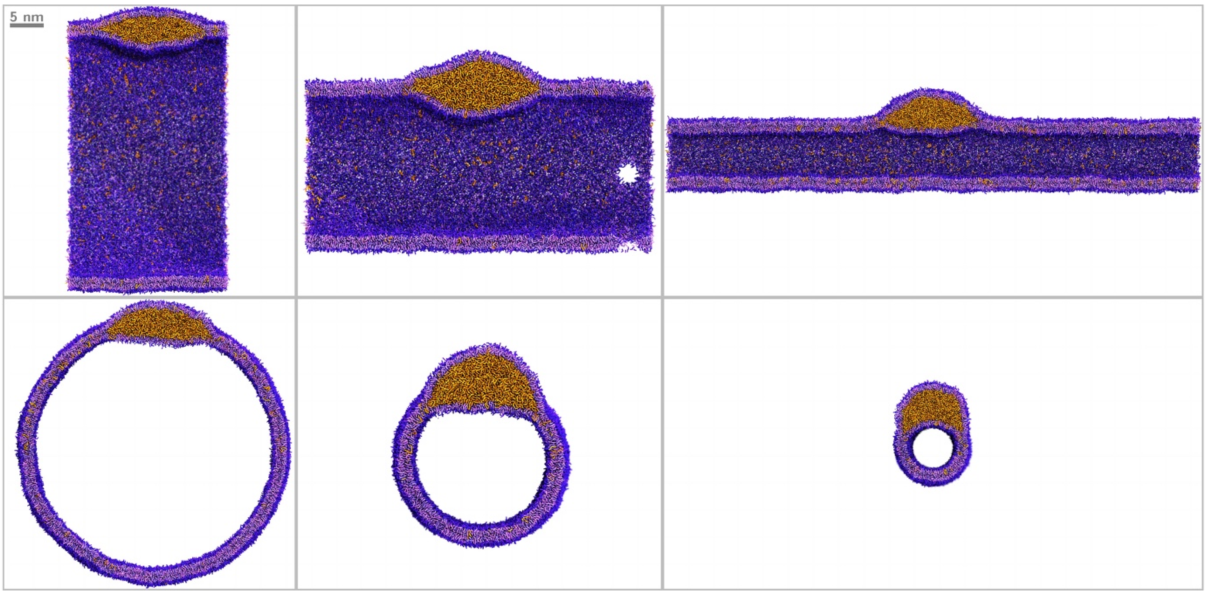

Figure S1. Nascent LDs embedded in membrane tubules (DOPC lipids) with a diameter of 10 nm, 35 nm, and 50 nm, generated by extending the simulation box. Free changes in tubule size are allowed by the presence of hydrophilic pores, kept open by an external potential.

##### 4. Membrane tubules with pores have negligible surface tension

Calculations of surface tension in large systems (both planar and tubular) are problematic due to wide structural fluctuations. Such calculations are even more complicated in the presence of nascent LDs, due to the lack of symmetry in the system. On the other hand, in tubular systems, it is simple to calculate the force acting along the tubule, as such force is related to the pressure, in the simulation box, in the direction of the tubule axis. Such pressure has an isotropic component due to bulk water, and a component due to the cylindrical membrane, that is curved. The (contractive) force due to the curvature energy of the membrane is related to the bending rigidity of the membrane[14]:

$$F_x = \frac{2\pi k_c}{R} \qquad \text{(eq. 1)}^{14}$$

where $k_c$ is the bending modulus of the membrane and *R* the radius of the cylinder. Note that, in an experimental setting, the radius of tubules pulled from vesicles is generally difficult to measure (the size is smaller than optical resolution), and is therefore calculated from the force (measured, for example, with optical tweezers, see ref. 15) and the tension (measured on a vesicle continuous with the tubule, by micropipette aspiration), using:

$$R = \sqrt{\frac{k_c}{2\sigma}} \qquad \text{(eq. 2)}^{16}$$

Notice that, based on eq. 1 and 2, the square of the force is linearly related to the mechanical tension in the system:

$$F_x = 2\pi\sqrt{2\sigma k_c} \qquad \text{(eq. 3)}^{16}$$

In simulations, the force can be calculated from the pressure in the system:

$$F_x = (P_{xx} - P_{bulk})A_{yz} \qquad \text{(eq. 4)}^{14}$$

where $P_{bulk}$ is the pressure due to water (1 bar) and $A_{yz}$ is the box *yz* area (normal to the direction of the tubule). From the force, we can also estimate the total mechanical tension σ acting on the tubular membrane:

$$\sigma = \frac{F_x}{4\pi R} \qquad \text{(eq. 5)}^{16}$$

Simulations of DOPC tubules with 1.5% TG (below the nucleation threshold, TG is well dispersed in the bilayer membrane) yielded a force of 22 pN when the tubule had a diameter of about 18 nm; this is similar to the force measured experimentally when pulling tethers from vesicles[15,16].

If the material is homogeneous and no additional forces act on the tubule, from $F_x$ we can calculate the bending modulus of the membrane.

$$k_c = \frac{F_x R}{2\pi} \qquad \text{(eq. 6)}^{14}$$

Tubules with smaller diameter give a stronger force and approximately the same $k_c$ (Table S2) – as expected for homogeneous systems. The result obtained from the cylinder is in good agreement with the one obtained from the buckling method[12].

The good match between bending moduli calculated with two different methods strongly supports that mechanical tension in the tubules originates from curvature energy, and interfacial tension (surface tension, in the main text) is negligible.

**Table S2.** Force along the direction of the tubule in equilibrium simulations, in the absence of LDs. Systems are homogeneous, so the force can be used to calculate the bending rigidity of the bilayer membrane.

| System | # PL | # TG | Simulation time (μs) | $L_x$ (nm) | $R$ (nm) | $P_{xx}$ (bar) | $P_{bulk}$ (bar) | $F_x$ (pN) | $k_c$ ($k_BT$) |
|---|---|---|---|---|---|---|---|---|---|
| DOPC tubule | 22482 | 350 | 20 | 75 | 18.4 | 0.938 ± 0.006 | 1.004 ± 0.006 | 22 ± 3 | 16.7 ± 3 |
| DOPC tubule | 22482 | 350 | 20 | 90 | 15.3 | 0.907 ± 0.005 | 1.013 ± 0.007 | 31.5 ± 2 | 18.5 ± 2 |

#### 5. Sorting of lipids in membrane tubules with embedded nascent LDs

A series of POP-MD simulations of LD growth was carried out by emulating TG synthesis at constant number of phospholipids, while keeping hydrophilic pores open. Some asymmetry in composition developed during the simulations and the subsequent equilibration runs. We analyzed differences in composition between inner and outer leaflet, as well as between bilayer and monolayer region, using a customized version of the SuAVE software[10,11].

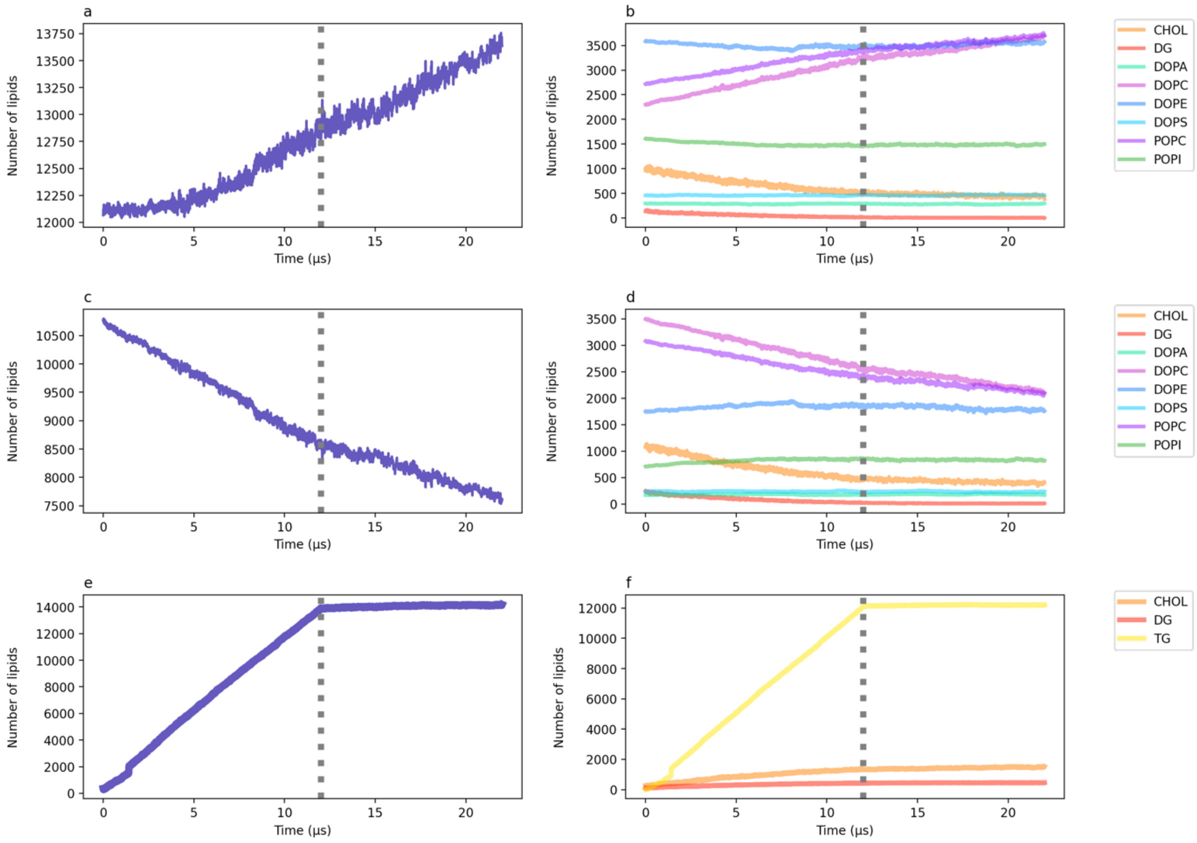

Figure S2. Leaflet asymmetry in LD growth simulations and subsequent equilibration, in tubules with the ER composition, in the absence of seipin; only TG is added to the system. The first 12 μs correspond to the growth phase (POP-MD), and the last 10 μs (indicated by the grey dotted line) correspond to the subsequent equilibrium simulation. (a) Total number of lipids and (b) lipid compositions in the outer leaflet. (c) Total number of lipids and (d) lipid compositions in the inner leaflet. (e) Total number of lipids and (f) lipid compositions in the LD core.

Hydrophilic pores allow growth in the overall number of lipids in the outer leaflet (Fig. S2a), but the growth is highly inhomogeneous: in fact, most phospholipids do not grow at all, while the number of DOPC and POPC shows a major increase (Fig. S2b). The inner leaflet shows the opposite picture: the overall number of lipids decreases (Fig. S2c), and this is due mostly to a decrease in the number of DOPC and POPC lipids (Fig. S2d). DG and cholesterol decrease in both the inner and outer leaflet, as they partially dissolve into the bulk oil phase (Fig. S2f). Notice that hydrophilic pores allow flip flop of all lipids, and sorting of phospholipids is due to their intrinsic preference for different regions in the system. Specifically, PC lipids prefer the positively curved outer surface, while PE and PI prefer the negatively curved inner leaflet.

#### 6. Comparison between Drosophila and yeast seipin structure

We used the seipin structure from Drosophila (PDB code 6MLU[17]), adding the transmembrane (TM) and the N-terminal helices based on predictions by AlphaFold2[18]. In the cryo-EM structure, the N-terminal helix (residues 20-56), the TM domains (residues 57-77 and 253-273), and two loops connecting the TM domain to the luminal domain (residues 78-88 and 242-252) are missing. We used Modeller[19] to build the complete atomistic structure. The electronic density map of 6MLU shows the presence of the TM domains and the N-terminal helix at lower density, suggesting that these parts of the protein are mobile. Therefore, in the Martini model, no elastic network was added to the TM domains and the N-terminal helix, to allow high mobility. An elastic network was applied to conserve the structure of the luminal domains (residues 89-241). Comparison with the structure of yeast seipin[20] shows high similarity of the luminal domain in the two structures (Fig. S3). When compared by TM-Align[21], the luminal domains yield a TM-score of 0.58, where a score between 0.5 and 1 indicates that the two proteins have approximately the same fold. The transmembrane helices, not shown in the figure, are of near-identical length.

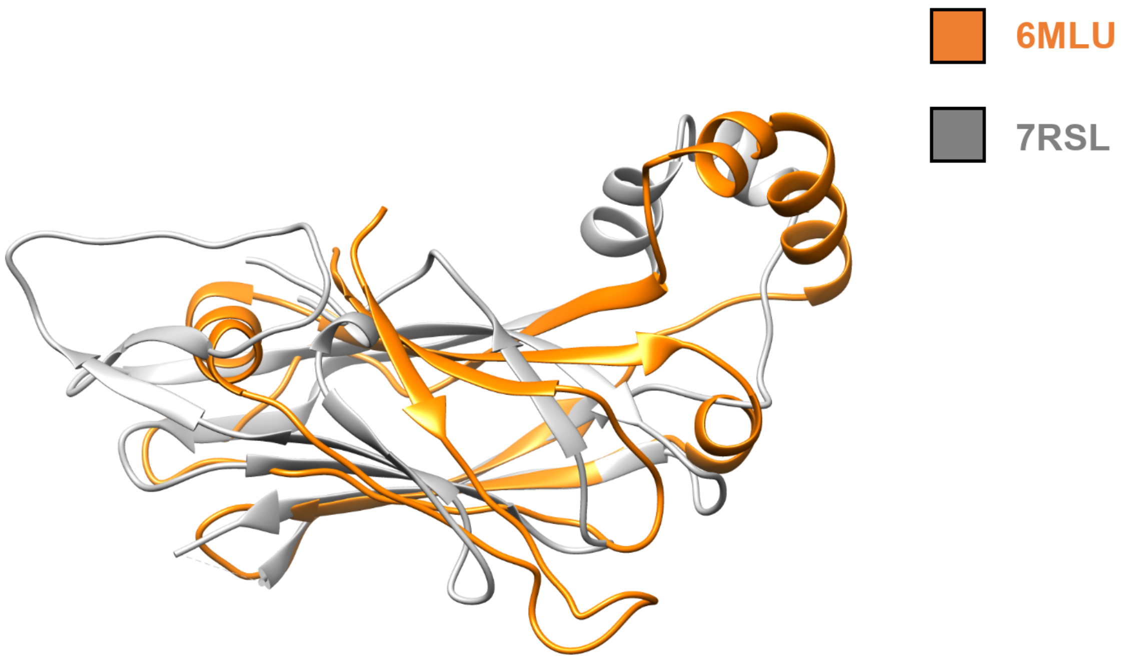

Figure S3. Comparing the luminal domains of Drosophila and yeast seipin structures (PDB codes 6MLU and 7RSL).

#### 7. Surface tension in tubular systems with nascent LDs

The theory summarized above (SI.4) is only applicable to homogeneous systems. As highlighted by Sorre et al., "departure of the square of the force, $F_x^2$ vs $\sigma$ (mechanical tension) from linearity reflects a difference between tube and mother vesicle compositions"[15]. In our case, the tubular membrane is not connected to a vesicle, but the nascent LD represents an inhomogeneity in the system, with different properties compared to the tubular membrane. In the presence of nascent LDs in the tubule, the amount of TG in the bilayer membrane remains approximately constant (1.5~2% of the phospholipid content with Martini v.3, about 3.5% in Martini v.2), hence the contribution from curvature energy of the tubule to the contractive force $F_x$ also remains approximately constant. However, the LD inclusion may induce changes in bending rigidity (probably decreasing, since the oily portion behaves like an isotropic liquid), and interfacial tension (probably increasing, due to the presence of an oil/phospholipid/water interface) in the non-bilayer region of the system. In simulations of LD growth, tension is difficult to assess not only because of large undulations, but also due to the non-equilibrium nature of POP-MD and the lack of symmetry in the system. On the other hand, the force exerted along the tubule can be calculated easily and precisely in equilibrium simulations, as done in the absence of nascent LD (SI.4).

We carried out simulations of tubules with embedded LDs of different sizes in equilibrium conditions (i.e., no addition of any molecule), starting from frames generated by POP-MD with *synthesis of TG only*. The force along the tubule ($F_x$) increased abruptly with LD nucleation (Table S3, orange shading). The value of $F_x$ obtained in the presence of a small nascent LD (2500 TG molecules) is about 3 times higher than observed for the tubule without LD. In the case of a larger LD (7500 TG molecules), the force is nearly an order of magnitude stronger, indicating a strong tendency of the system to contract along the tubule axis. Considering the structural inhomogeneity, calculations of membrane bending rigidity (eq. 6) or mechanical tension (eq. 5) would be meaningless. We claim that the very high contractive force along the tubule axis is due to an increase in interfacial tension in the system. Direct calculation of mechanical tension in the system (via the local stress tensor) is very challenging, therefore we cannot say if tension increases only in the monolayer region or also in the bilayer. However, *if the increase in the force is indeed due to surface tension, then an increase in phospholipid density during LD biogenesis must restore a low force during LD nucleation and growth.*

To verify our hypothesis, we repeated the equilibrium force calculations (Table S3, grey shading) starting from POP-MD snapshots taken from simulations with *simultaneous synthesis of TG and PL*, with PL added to both leaflets. The systems are very similar to the ones described above, as they contain one nascent LD of relatively small size; the only difference is that PL were added to the tubule together with TG (PL:TG ratio 1:1), hence the tubule (which has fixed length) contains more PL. In this case, remarkably, the force along the tubule is nearly identical to the force obtained in the absence of nascent LD. This indicates that, in simulations, phospholipid synthesis is necessary to maintain low surface tension in the system.

**Table S3.** Pressure along the direction of the tubule in equilibrium simulations, in the presence of nascent LD or fully budded LD.

| System | # PL | # TG | Simulation time (μs) | $L_x$ (nm) | $R$ (nm) | $P_{xx}$ (bar) | $P_{bulk}$ (bar) | $F_x$ (pN) |
|---|---|---|---|---|---|---|---|---|
| DOPC tubule (no LD) | 22482 | 350 | 20 | 75 | 18.4 | 0.938 ± 0.006 | 1.004 ± 0.006 | 22± 3 |
| DOPC tubule + nascent LD (POP-MD: TG only) | 22482 | 2500 | 20 | 75 | 18.3 | 0.83 ± 0.009 | 0.996 ± 0.008 | 60 ± 5 |
| DOPC tubule + nascent LD (POP-MD: TG only) | 22482 | 7725 | 20 | 75 | 18.5 | 0.57 ± 0.01 | 0.999 ± 0.009 | 160 ± 5 |
| DOPC tubule + nascent LD (POP-MD: TG + DOPC) | 23982 | 1850 | 20 | 75 | 18.9 | 0.939 ± 0.01 | 1.00 ± 0.01 | 22 ± 4 |
| DOPC tubule + nascent LD (POP-MD: TG + DOPC) | 25482 | 3350 | 20 | 75 | 19.2 | 0.908 ± 0.01 | 0.97 ± 0.01 | 23 ± 4 |

#### 8. Asymmetric phospholipid synthesis is necessary and sufficient for LD budding

We repeated simulations of LD growth by simulating asymmetric synthesis of PL with POP-MD using different LD sizes (from ca. 600 to over 7000 TG molecules), different chemical composition of the membrane (pure DOPC, ER mixture, and ER mixture enriched with lyso-PC), and different membrane geometry (flat vs. tubular), see Table 1. *Asymmetric PL synthesis induced budding of LDs towards the cytosolic compartment in all cases*, independently of LD size, membrane composition, presence or absence of lysolipids, and membrane geometry. The budding transition produced structures in which the LD volume was entirely on one side of the bilayer membrane, which allows us to propose a working definition of the budding transition. *The budding mechanism was indistinguishable in all cases*: the onset of the budding transition was observed at approximately the same degree of leaflet imbalance, i.e., the kinetics of budding was about the same in all simulations. Even the addition of 10% lyso-PC (replacing 5% DOPC and 5% POPC in the ER mixture) did not result in any significant difference (Fig. S4b-c, S4d-e). We conclude that LD volume, the nature of the phospholipids, and membrane geometry make no substantial difference to the budding mechanism; once TG synthesis yielded a nascent LD, asymmetric phospholipid synthesis is the main driving force for LD budding.

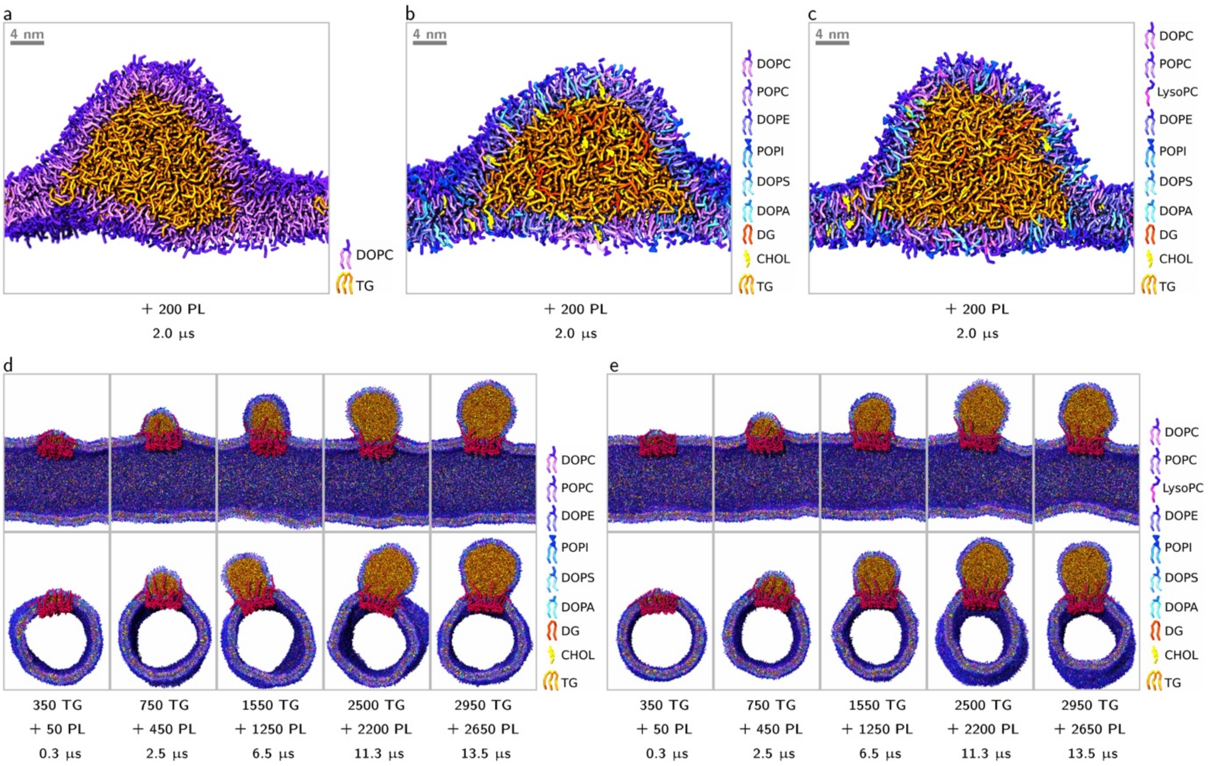

Figure S4. POP-MD simulations of flat bilayer membranes consisting of 2016 PL and 625 TG; the number of TG molecules in the nascent LD was less than 600 in all cases; the membrane contained (a) only DOPC, (b) the ER mixture, and (c) the ER mixture enriched with lyso-PC. (d) POP-MD simulations of tubular bilayer membranes (23182 PL and 300 TG before POP-MD) using the same ER mixture, and (e) the ER mixture enriched with lyso-PC.

#### 9. Formation of defects in the LD neck and in tubules

We used POP-MD to simulate the synthesis of TG and PL in flat bilayer membranes, in systems initially containing a nascent LD and no seipin, inserting TG in the proximity of the existing nascent LD and PL in random locations. Monolayer-bounded defects formed at the LD neck, exactly as in simulations of membrane tubules (Fig. S5a). Both in tubular and in flat membranes, defects are caused by the excess phospholipid surface density of the outer leaflet: when the surface density of phospholipids is over a certain threshold, neither the bilayer nor the monolayer can accommodate more phospholipids on their surface, therefore the area of the interface increases. Remarkably, defects are monolayer-bounded, similar to the ones formed by the collapse of lipid monolayers at the air-water interface[22].

In contrast, in the presence of seipin, excess phospholipids caused defects away from the LD neck, in the membrane tubule (Fig. S5d). Different from defects in the LD neck, the ones in the tubule were bilayer-bounded.

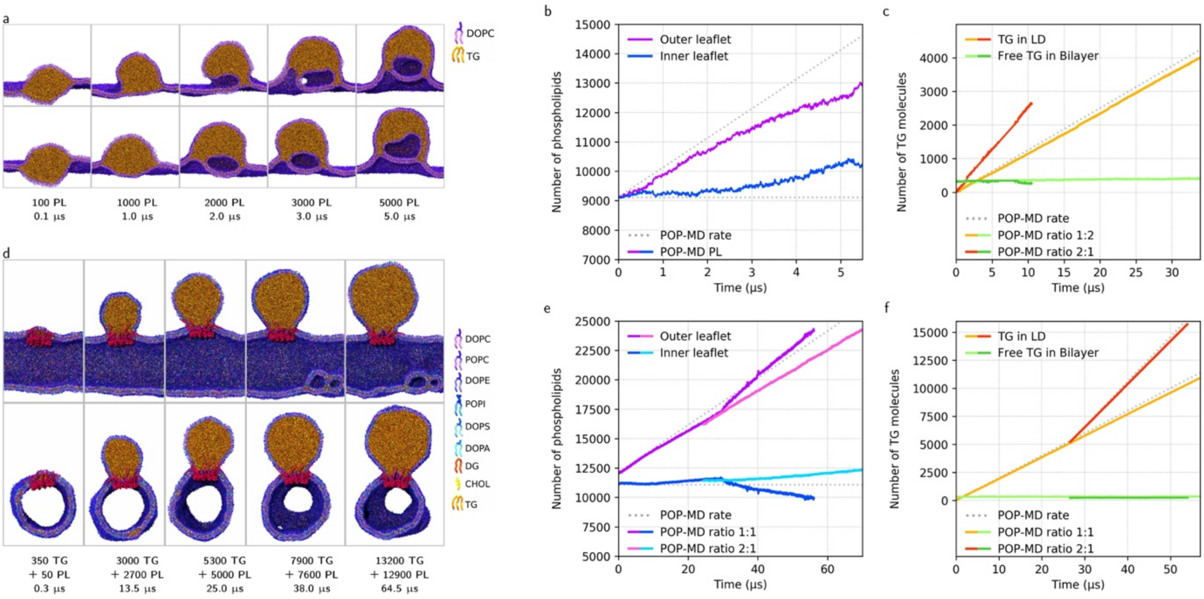

Figure S5. (a) Formation of water-filled defects in flat bilayer membranes (18144 PL and 7500 TG molecules before POP-MD), at the LD neck, protruding into the oil region. (b) Number of phospholipids in the inner and outer leaflet as a function of time. (c) Number of TG molecules in the LD and dissolved in the phospholipid bilayer. (d) Formation of lipid-filled defects in tubular membranes, away from the LD neck, protruding into the lumen of the tubule. (e) Number of phospholipids in the inner and outer leaflet as a function of time. (f) Number of TG molecules in the LD and dissolved in the phospholipid bilayer.

#### 10. Seipin minimizes the number of nucleated LDs by localizing TG synthesis

In the absence of seipin, synthesis of TG and PL leads to multiple nucleation and multiple budding events. This is consistent with the observation that mutations or depletion of seipin induce aberrant phenotypes, with few giant LDs and numerous tiny LDs[23-25]. Previous simulations suggested that seipin reduces the number of nucleated LDs due to the high affinity of seipin TM helices and $\alpha$2-$\alpha$3 helix (in luminal domain) for TG, preventing TG from diffusing out of the oligomeric protein ring[26,27]. The $\alpha$2-$\alpha$3 helices are necessary for seipin functioning in forming TG clusters. However, such simulations were carried out in equilibrium conditions, with a fixed number of TG molecules, while TG synthesis brings the system out of equilibrium, and may generate inhomogeneous distributions. To better understand how seipin reduces the number of nucleated LDs, we simulated the synthesis of TG molecules (a) in the proximity of the seipin oligomer, (b) away from seipin, or (c) in random positions in the membrane.

Localization of TG synthesis has a major effect on LD nucleation: when TG was synthesized close to the seipin ring, one nascent LD nucleated within the ring; on the other hand, when TG was synthesized far from seipin or in random positions, multiple nascent LDs nucleated at the same time, in and out of the seipin ring, both in tubular and in flat membranes (Fig. S6). This is explained by the strong tendency of TG to phase separate from phospholipids and the relatively long diffusion times required for TG molecules to travel within the tubular membrane to the seipin ring, due to the large size of the system and the limited time scale of the simulations. Extending the simulations, multiple LDs coalesced, and the one scaffolded by seipin showed higher stability, as expected. However, even on the relatively short length scale of our flat bilayer membranes (78x78 nm) and tubules (75 nm), coalescence of all LDs into one was not observed on the simulation time scale (10-15 $\mu$s). It can be argued that multiple nascent LDs are observed in the simulations due to the relatively short simulations times, insufficient for diffusion of the nascent LD. However, the surface area of ER tubular region in cells is much higher than in the simulated tubular membrane, implying much longer diffusion times for LDs nucleated far from seipin. We suggest that, in cells, seipin reduces the number of nucleating LDs not only by catalyzing TG nucleation within its scaffold, but also by localizing TG synthesis, therefore reducing the time required for TG diffusion. This is compatible with experimental data indicating an interaction between seipin and TG-synthesizing enzymes[28,29]. We propose that synthesis of TG away from seipin would cause nucleation of multiple nascent LDs, leading to the aberrant phenotypes described in the literature[24,30,31].

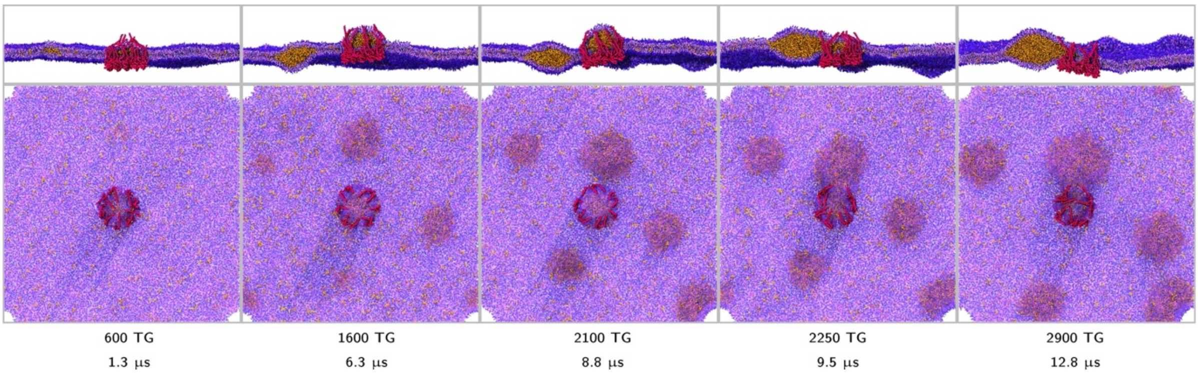

Figure S6. Effect of localization of TG synthesis during LD growth, in the presence of seipin. Inserting TG in random position in a flat membrane, in the presence of seipin, produces multiple nucleated LDs, in and out of the seipin ring.

##### 11. The localization of phospholipids synthesis does not affect LD budding mechanism

We repeated the simulations of LD growth in the tubular membranes mimicking the complex ER composition, in the presence of seipin, by adding oil and phospholipids in a 1:1 ratio, in 3 different ways: phospholipids were added in the proximity of the seipin ring (Fig. S7a), on the opposite side of the seipin ring (Fig. S7b), or in random locations of the outer leaflet (Fig. S7c). In all three cases, LD budding followed the same mechanism, resulting in a regular cylindrical shape of the ER tubule, with no deformations in the LD monolayer nor close to the LD neck.

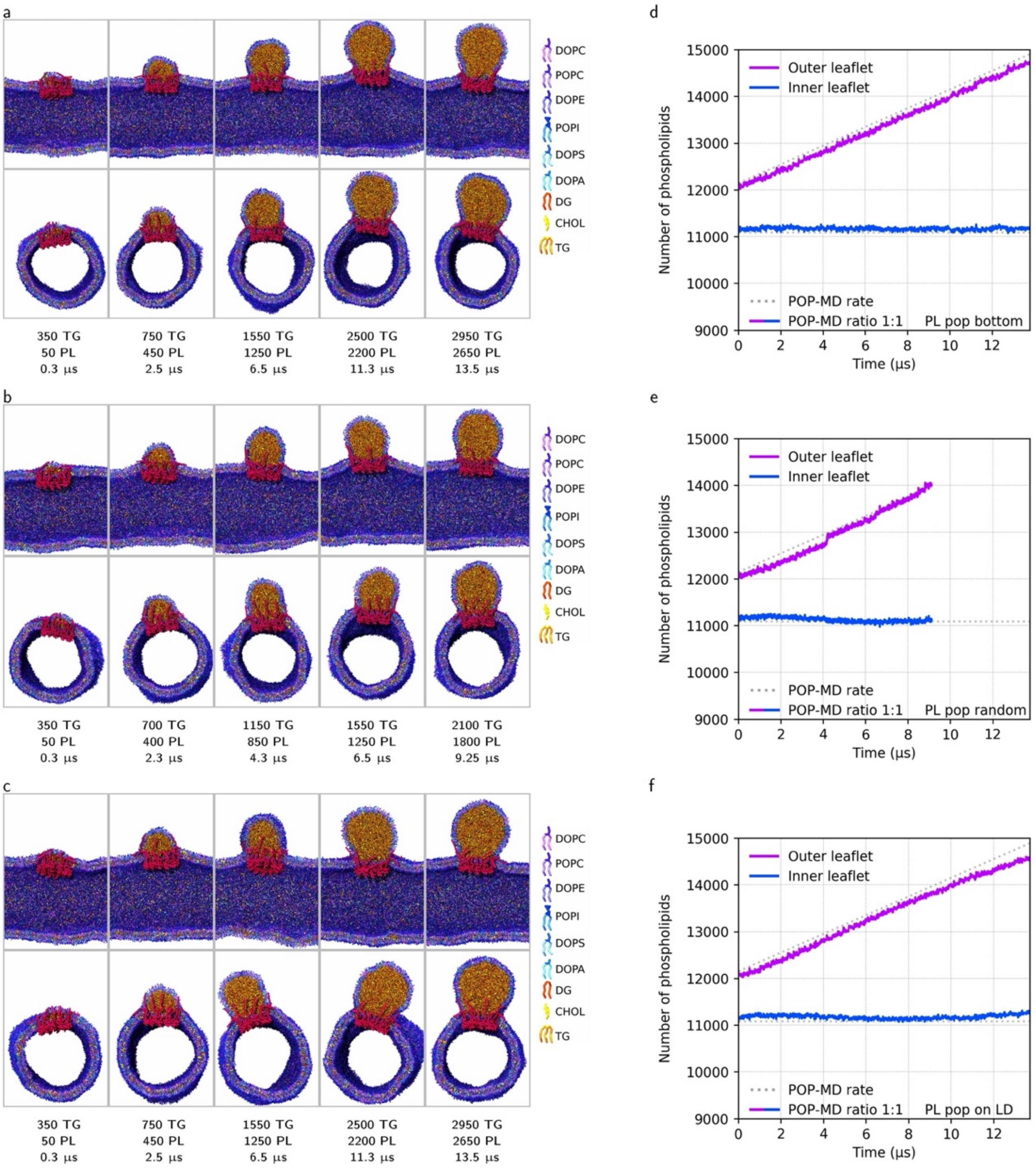

Figure S7. Growing a TG droplet in a tubular membrane, in the presence of seipin, by synthesis of PL in different regions of the tubular membrane, using a complex ER mixture (8 different PL types). PL molecules are inserted in the outer leaflet (a) at the bottom of the tubular membrane, (b) in random locations of the tubular membrane, and (c) close to the nascent LD. Build-up of leaflet asymmetry in the corresponding simulations, (d), (e), and (f).

**12. Sorting of lipids between inner leaflet, outer leaflet, and monolayer upon LD budding**

We analyzed sorting of different lipids during LD growth and budding in tubules containing the complex ER-mimicking lipid mixture (composition described above). Hydrophilic pores were present throughout the simulation, and PL were synthesized only in the outer leaflet. Although the use of hydrophilic pores may appear to defeat the purpose of asymmetric PL synthesis, asymmetry between the two membrane leaflets persisted during the simulation, both in the non-equilibrium growth phase (Fig. S8, 0-70 μs) and during the subsequent equilibration phase (Fig. S8, 70-81 μs).

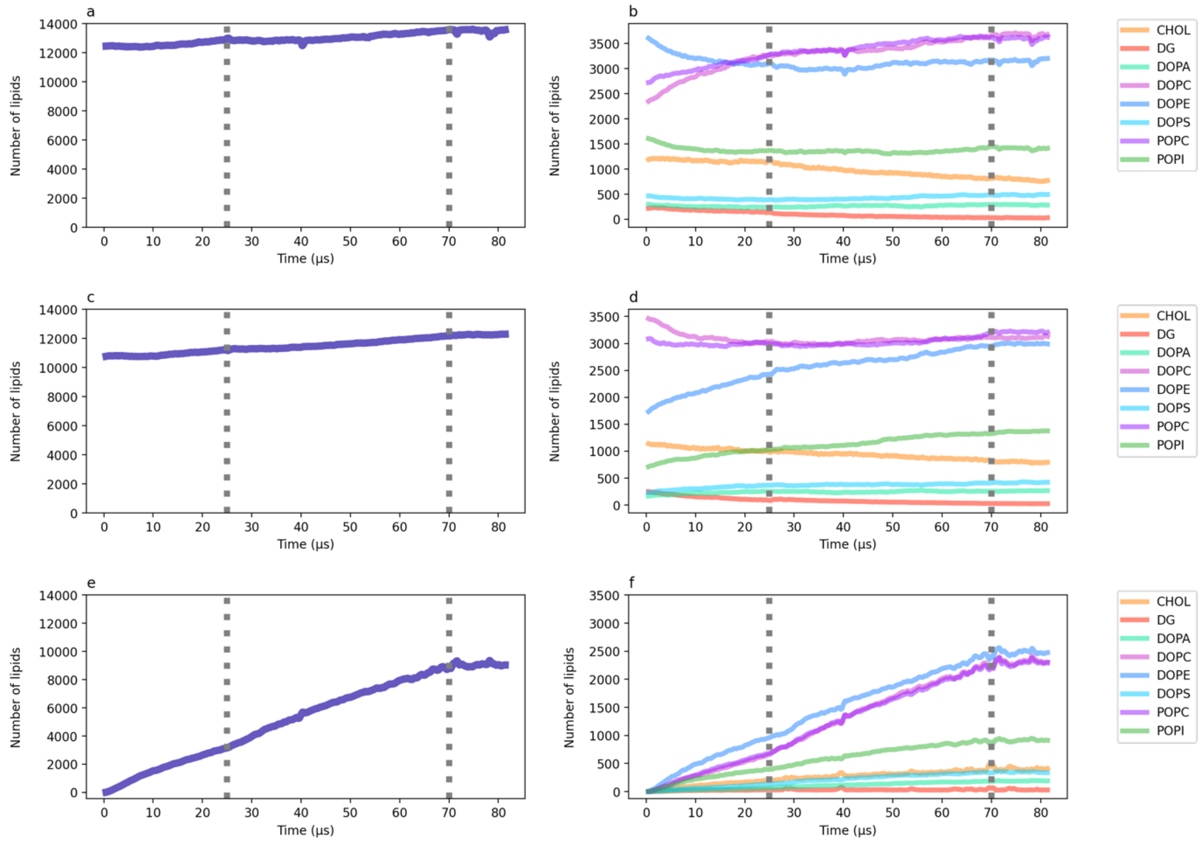

Fig. S8. Leaflet asymmetry in LD growth simulations and subsequent equilibration, in tubules with the ER composition, in the presence of seipin; both TG and phospholipids are inserted in the system by POP-MD. The first 25 μs correspond to a 1:1 PL:TO synthesis rate; the subsequent 45 μs (i.e., 25-70 μs) correspond to a 1:2 PL:TO synthesis rate, separated by a grey dotted line. The second dotted line marks the end of the POP-MD, and is followed by equilibrium MD (70-81 μs). (a) Total number of lipids and (b) individual lipid compositions in the outer tubule leaflet. (c) Total number of lipids and (d) individual lipid compositions in the inner tubule leaflet. (e) Total number of lipids and (f) individual lipid compositions in the PL monolayer surrounding the LD core.

For each portion of the system, the lipid composition changes significantly during non-equilibrium POP-MD, but not during the equilibrium MD phase, suggesting that the synthesis rate is slow enough to produce systems not very far from thermodynamic equilibrium in terms of lipid distribution.

During the LD growth phase (0-70 μs), we observed a small but steady growth in the overall number of lipids in both leaflets overall, alongside with a minor increase in the diameter of the tubule; we also observe a large and steady increase in the overall number of lipids in the monolayer bounding the budded LD, due to the increase in LD volume (due to TG synthesis).

Moreover, as the LD grows, the amount of cholesterol (CHOL) and dioleoylglycerol (DG) decreases in all interfacial regions (outer leaflet, inner leaflets, and LD monolayer). The decrease in CHOL and DG occurs despite all lipids are inserted in the outer leaflet, and is explained by partitioning of both CHOL and DG (to a limited extent) into the LD core. Partial redistribution of CHOL and DG to the LD core was also observed in LD growth simulations without addition of phospholipids (Fig. S2f). Since part of CHOL and DG are not found at the interface, we calculated an "adjusted" fraction of each phospholipid that does not take into account CHOL and DG (Table S4).

Table S4. Fraction of phospholipids, averaged over the last 11 μs of simulation (equilibrium MD, no phospholipid synthesis). The adjusted fraction of each lipid is calculated by excluding CHOL and DG from the count.

| **Lipid** | **overall lipid fraction** | **adjusted overall lipid fraction** | **lipid fraction in the outer leaflet** | **lipid fraction in the inner leaflet** | **lipid fraction in the monolayer** |
|---|---|---|---|---|---|
| **DOPC** | 0.250 | **0.284** | 0.289 | 0.272 | 0.269 |
| **POPC** | 0.250 | **0.284** | 0.283 | 0.281 | 0.266 |
| **DOPE** | 0.230 | **0.261** | 0.249 | 0.262 | 0.288 |
| **DOPS** | 0.030 | **0.034** | 0.038 | 0.036 | 0.040 |
| **DOPA** | 0.020 | **0.023** | 0.023 | 0.023 | 0.022 |
| **POPI** | 0.100 | **0.113** | 0.111 | 0.119 | 0.106 |

As the LD grows, inhomogeneities in the composition of the system develop, and persist throughout the equilibrium phase (70-81 μs). In particular, the composition of the outer and inner leaflets is not the same: the outer leaflet is enriched in DOPC and POPC and depleted in DOPE, while the reverse is true for the inner leaflet. Remarkably, the composition of the LD monolayer is different from the composition of the outer leaflet: DOPE is significantly enriched (Fig. S8f and Table S4), despite its negative intrinsic curvature; at the same time, both DOPC and POPC are depleted, and the fraction of cholesterol at the interface is significantly higher than in each bilayer leaflet. Outer leaflet and LD monolayer are contiguous, and differences in their respective composition indicate that simulations can reproduce the different chemical potential of each lipid in different regions of the system, also observed experimentally[32].

#### 13. LD-bilayer connection is extremely stable

Having simulated LD budding under different conditions, we never observed mechanical instability of the LD neck nor any tendency of the budded LD to detach from the membrane tubule, not even when the LD neck was highly distorted by water-filled defects. High stability of the LD neck is compatible with experimental results, showing that mature, fully budded LDs generally remain in contact with the ER via an LD-ER contact site[33] (usually punctuated by seipin).

To further probe the stability of the LD-tubule connection, we stressed the system by applying external mechanical forces via steered MD simulations[34]. To this end, we simulated a vesicle with an embedded LD and added a strong external force pulling the LD away from the vesicle at constant speed (5 nm/µs). After pulling for 10 µs, the center of mass of the TG LD was displaced by 50 nm away from the center of mass of the vesicle. During pulling, the LD neck became longer and thinner, reaching an outer diameter of only 9.5 nm (Fig. S9, 12 µs), very close to the thickness of two bilayer membranes. Despite the extremely small diameter, the tubular neck did not rupture. Extension of the simulation for 10 µs with forces applied in fixed position also did not produce rupture (Fig S9, 22 µs). Release of external forces resulted in recovery of the original shape in approximately 20 µs: the neck becomes shorter and wider, reaching a final diameter of 30 nm (same as the initial diameter). Analogous simulations with external pulling forces were also carried out on tubular membranes, with higher pulling rates, and confirmed the high mechanical stability of the LD-ER connection.

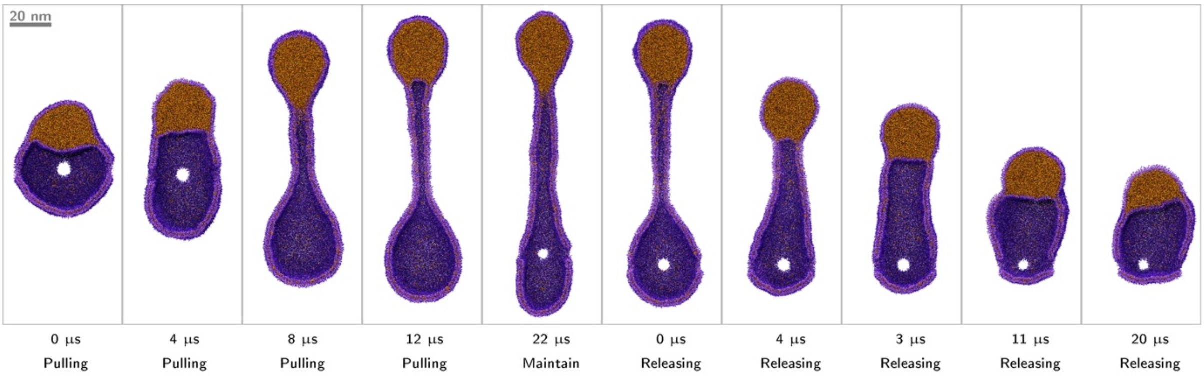

Figure S9: Pulling and releasing of a nascent LD in a vesicle: the LD is pulled during the initial 12 µs of steered MD, then it is held in place for 10 µs, and finally forces are released during a 20 µs relaxation run. The presence of hydrophilic pores does not affect the stability of the LD-bilayer connection.

#### 14. Spreading of surface density and surface tension in bilayer systems

We used POP-MD to increase the surface density of lipids in bilayer systems consisting of pure DOPC, with linear sizes of 30-40 nm. Lipids were added only to half of the bilayer area, and we calculated the time required for the two halves to reach the same surface density – corresponding to the same (vanishing) surface tension. After insertion of phospholipids, surface density becomes homogeneous within a few ns, indicating that, in bilayer systems, surface tension propagates with speed of the order of m/s (Fig. S10). These results are consistent with experiments showing that the energy to move lipids from one region to another of a fluid membrane is small, and tension-driven membrane dynamics is fast[35-37].

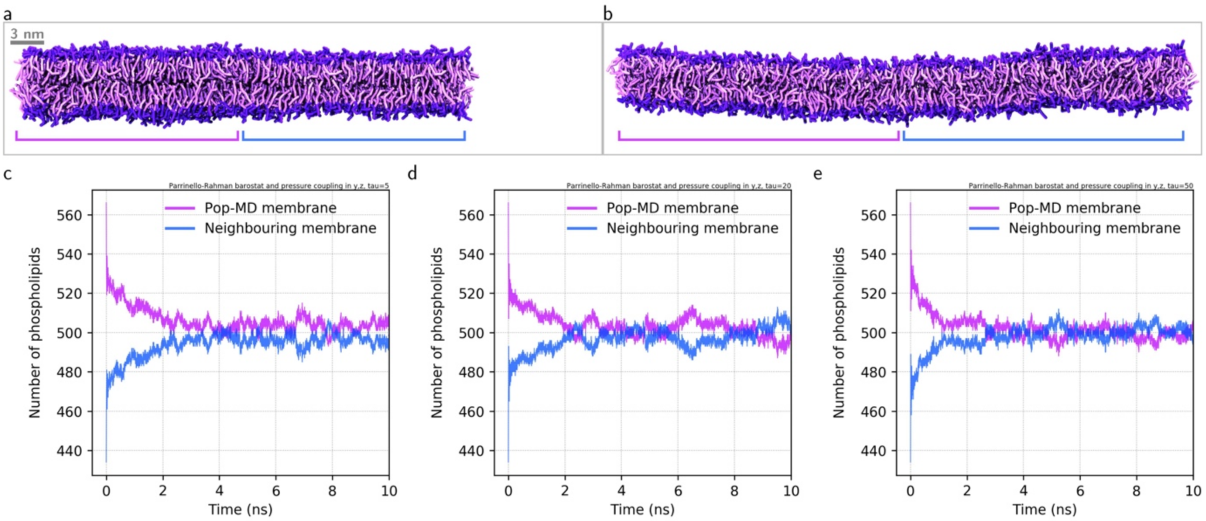

Figure S10. Spreading of surface density in simulations of flat bilayer membrane (100% DOPC) with inhomogeneous lipid distribution. Inhomogeneity was generated by inserting phospholipids only in the left-hand side of the bilayer; snapshot at the moment of insertion (a) and after 10 ns (b). (c) Number of phospholipids in the bilayer membrane after lipid insertion (at time zero); the number of phospholipids in each half of the membrane, the one with the inserted lipids (purple line) and the other (blue line), converges in a few ns. Simulations were carried out with the Parrinello-Rahman barostat[8] using different parameters for the coupling constant $\tau$: $\tau$ =5 ps (c), $\tau$ =20 ps (c), and $\tau$ =50 ps (e).
